# LRRC57/RABIN is a presynaptic inhibitor of Rab GTPases in glutamatergic neurons

**DOI:** 10.64898/2026.08.05.743108

**Authors:** Dušan Garić, Jonathan G. Murphy, Brett J.W. Teubner, Christopher M. Davenport, Chablis D. Shreffler, Laura J. Janke, Woo Jung Cho, Zhiping Wu, Meghan McReynolds, Sarayut Phasuk, Jason Y. Fu, De’Keerah A. Lewis, Thomas Confer, Jonathon Klein, Melissa Johnson, Camenzind G. Robinson, Shondra M. Pruett-Miller, Richard J. Heath, Junmin Peng, Jason D. Vevea, Stanislav S. Zakharenko

## Abstract

Synaptic vesicle cycling, if not properly constrained, can result in excessive neurotransmitter release and subsequent neural pathology. Rab GTPases orchestrate synaptic vesicle trafficking through GTP-dependent interactions with effector proteins, but the restraining mechanism of these interactions is unknown. Here we identify LRRC57 (or RABIN for RAB INhibitor), a conserved brain-enriched protein in glutamatergic synapses that binds multiple GTP-loaded synaptic Rabs and competitively blocks access to their effectors. Loss of *Rabin* increased glutamate release, expanded vesicle pools, accelerated vesicle turnover, and produced circuit hyperexcitability with epileptiform activity, which was mitigated by an antiepileptic agent that targets presynaptic function. Conversely, overexpression of the *Rabin* gene suppressed neurotransmitter release and protected against induced seizures and persistent epileptiform discharges. Together, these findings define a noncanonical decoy–effector mechanism that constrains presynaptic Rab signaling to preserve excitatory circuit stability.

## INTRODUCTION

The tightly coordinated process of synaptic vesicle (SV) exocytosis drives synaptic transmission. Synaptic transmission begins with the well-characterized, calcium-triggered fusion of SVs with the plasma membrane, a crucial step in neural function^1^. In contrast, the signaling networks that prepare SVs for exocytosis and constrain their release probability remain less well understood. Because synapses operate near a threshold for runaway excitation^2^, presynaptic mechanisms that actively limit vesicle release are essential for circuit stability, and their disruption is a common feature of epileptic and neurodegenerative disorders.

Rabs are small GTPases that act as molecular switches; they are active in their GTP-bound state and inactive after hydrolysis to GDP^3–7^. In their active form, Rabs regulate membrane trafficking across cell types, including vesicle budding, transport, tethering, and fusion. Each Rab occupies a distinct membrane domain and recruits specific effectors in a GTP-dependent manner, yet many of the more than 60 Rab family members still lack identified effectors^8^. Rab activity is classically regulated by guanine nucleotide–exchange factors (GEFs), GTPase-activating proteins (GAPs), and GDP-dissociation inhibitors (GDIs), which together determine Rab activation state, membrane association, and signal termination^6,7^. In contrast, Rab effectors are thought to transduce Rab signaling rather than directly limit or restrain it^3^.

At synapses, multiple Rab modules operate across the SV lifecycle^5,9^. SV-localized RAB3A-D (and possibly RAB27A/B) facilitate SV docking and release by interacting with the active zone complex, including the RAB3 effector RIM1α^10–14^. Additional Rab proteins contribute to vesicle biogenesis, transport, and recycling, suggesting that coordinated Rab signaling, rather than a single Rab pathway, governs vesicle release^5,15^. Disruption of Rab signaling is associated with epilepsy and neurodegeneration, underscoring the importance of precisely constraining Rab activity at nerve terminals^16,17^. However, whether Rab signaling at synapses is subject to inhibitory regulation beyond canonical GAP-mediated inactivation remains unknown. Current models imply that the local restraint of SV release depends on unknown molecular regulators^18,19^. However, the identity of such regulators and how they restrain SV release are unknown.

By interrogating the uncharacterized fraction of the brain proteome through an unbiased bioinformatic analysis of highly evolutionarily conserved proteins (>75% similarity between vertebrate and invertebrate Metazoan species), whose function is still unknown and whose expression is enriched in the brain, we identified *Lrrc57* (*ZK546.2* in *Caenorhabditis elegans* and *CG3040* in *Drosophila melanogaster*). *Lrrc57* displays a very high degree of evolutionary conservation across all Metazoa, indicating strong evolutionary pressure to preserve its function. In large-scale invertebrate screens, *Lrrc57* expression was neuron-enriched; knockdown caused partial lethality and behavioral defects in *C. elegans*^20,21^ and *D. melanogaster*^22^.

*Lrrc57* encodes a 27-kDa, horseshoe-shaped protein that consists of eight leucine-rich repeats; these repeats form a concave surface flanked by two highly conserved α-helices at the N and C terminus (**Fig. 1a**). This architecture does not confer any known catalytic activity but is present in various proteins of unrelated function, where it provides a versatile structural framework for the formation of protein–protein interactions. LRRC57 protein is N-myristoylated, and through its lipid tail, it is predicted to be associated with intracellular membranes^23,24^.

**Fig. 1.**
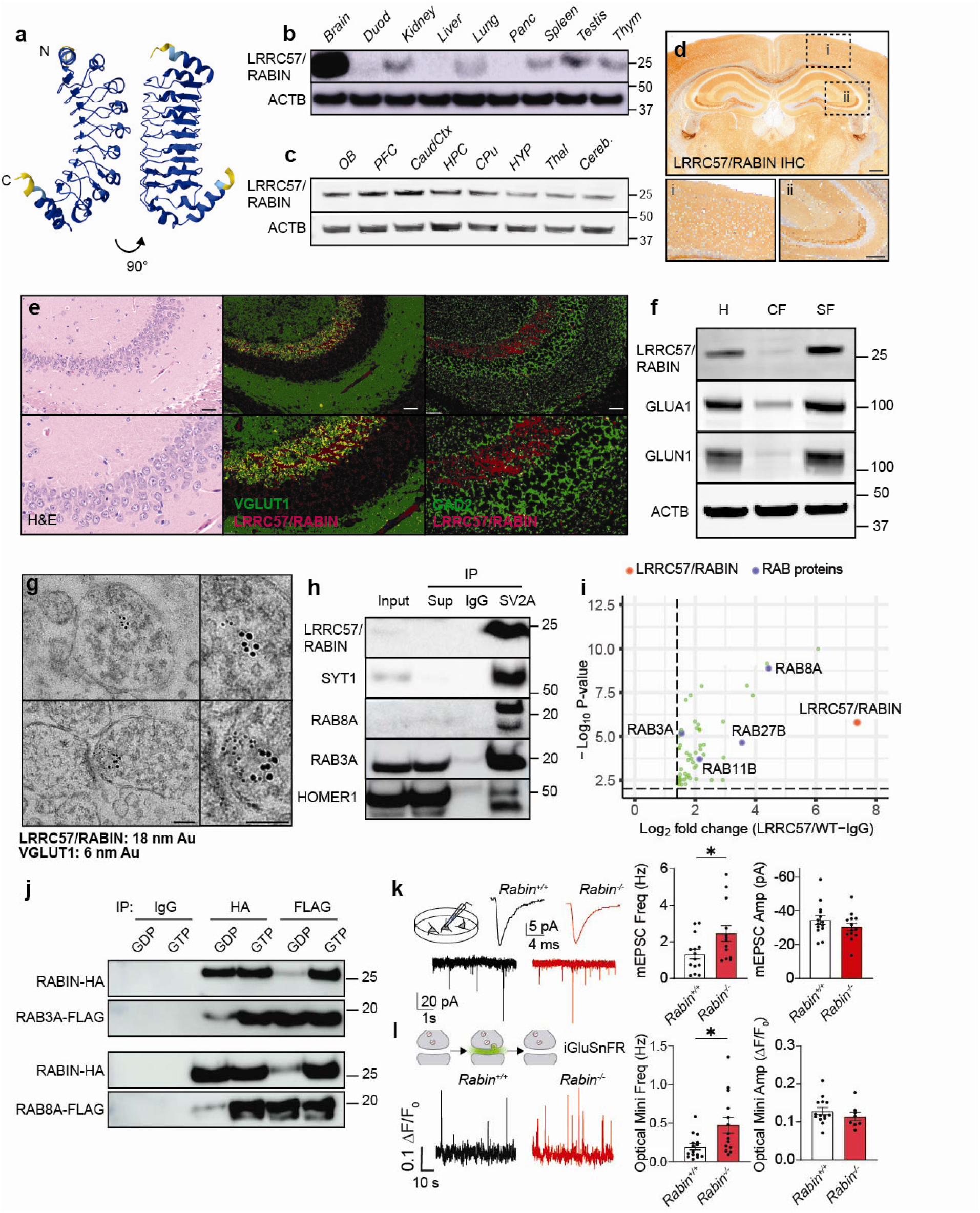
LRRC57/RABIN is localized to glutamatergic SVs and binds active Rab GTPases. **a.** Ribbon diagrams showing two views of the LRRC57/RABIN AlphaFold structural prediction. Color coding represents AlphaFold model confidence: blue, ≥90%, teal, 70%-90%, yellow, 50%-70%, orange, ≤50%. **b.** LRRC57/RABIN protein expression is enriched in the murine brain, relative to selected organs. **c.** LRRC57/RABIN protein expression is enriched in forebrain structures relative to other brain regions. **d.** Immunohistochemical localization of LRRC57/RABIN. Scale bar, 500 μm. Insets, LRRC57/RABIN expression in the cerebral cortex (i) and the hippocampus (ii). Scale bar, 250 μm. **e.** LRRC57/RABIN colocalizes with markers of excitatory but not inhibitory neurons. (Top panels) H&E staining shows immunostaining in the CA3 region. Co-immunolabeling of LRRC57/RABIN with VGLUT1 (excitatory neuron marker) and GAD2 (inhibitory neuron marker). Scale bars, 50 μm. (Bottom panels) 2× magnified insets. All immunofluorescence images are contrast-enhanced identically across conditions; segmentation was used only for display. **f.** Representative Western blots of mouse hippocampus homogenate (H) separated into a cytosolic fraction (CF) and a synaptoneurosome fraction (SF). **g.** Representative RABIN/VGLUT1 double-immunogold electron microscopic images of hippocampal CA3 synapses. Scale bars, 100 nm. **h.** Western blot analysis of synaptic vesicle (SV) proteins enriched by SV glycoprotein 2A (SV2A) co-immunoprecipitation. **i.** Volcano plot of the LRRC57/RABIN protein interactome identified by LRRC57/RABIN immunoprecipitation and proteomics of co-eluted proteins by mass spectrometry. Data were normalized against immunoglobulin (IgG) control samples. **j.** Representative Western blot depicts the GTP-dependence of RABIN–RAB binding in primary hippocampal neurons in culture. **k.** Representative recordings of miniature excitatory postsynaptic currents (mEPSCs) in primary hippocampal neurons harvested from embryonic day 18 (E18) *Rabin^+/+^* and *Rabin^−/−^*mice. Averaged single mEPSCs (above) and longer timescale traces (below) are shown. Frequency: Mann-Whitney *U* test, *U*=50, \**p*=0.048. n=14, 13 cells. Amplitude: Mann-Whitney *U* test, *U*=68, \**p*=0.280. n=14, 13 cells. **l.** Representative iGluSnFr fluorescence measured at individual synapses. Mean optical mini frequency was calculated across synapses within each field of view (FOV). Frequency: Welch’s two-tailed *t*-test, *t*=2.593, \**p*=0.0191, n=242, 233 synapses (15, 14 FOVs). Amplitude: Welch’s two-tailed *t*-test, *t*=0.9833, *p*=0.0.3377, n=231, 192 synapses (14, 9 FOVs). Data are presented as the mean ± SEM in **k** and **l**. **Abbreviations:** ACTB, β-actin; CaudCtx, caudal cortex; cereb, cerebellum; CPu, Caudate putamen; Duod, duodenum; H&E, hematoxylin and eosin; HPC, hippocampus; HYP, hypothalamus; IHC, immunohistochemistry; IP, immunoprecipitation; OB, olfactory bulb; Panc, pancreas; PFC, prefrontal cortex; Thal, thalamus; Thym, thymus; WT, wild type

Here we show that a family of well-known Rab GTPases (RAB3A, RAB8A, RAB11B, and RAB27B), which mediate the biogenesis, transport, and exocytosis of SVs, are the main interactors of the LRRC57 protein. LRRC57 localizes to glutamatergic SVs and preferentially binds the GTP-loaded forms of RAB3A and RAB8A, which is consistent with Rab effector–like behavior, and inhibits the interactions of these RABs with their respective effectors. Genetic disruption of *Lrrc57* resulted in excessive synaptic transmission, circuit hyperexcitability, and seizure susceptibility rescued by the antiepileptic levetiracetam, and increasing LRRC57 levels mitigated pathologic features of an epilepsy model. Collectively, our findings identify LRRC57 (or RABIN, for RAB INhibitor) as a previously unrecognized inhibitory regulator of Rab GTPases that constrains SV release and maintains neural circuit stability.

## RESULTS

### RABIN is localized to glutamatergic synaptic vesicles and binds active Rab GTPases

A systematic prioritization search for highly conserved, brain-enriched genes of unknown function identified *Lrrc57*/*Rabin* (**Figs. S1, S2a, Table S1**). To determine the expression and localization of RABIN, we developed a rabbit monoclonal antibody against a conserved C-terminal 12–amino acid sequence (**Fig. S2a**). Antibody specificity was validated by Western blotting and immunohistochemistry using newly generated conditional knockout (cKO) *Emx1^Cre^;Rabin^fl/fl^* mice (**Figs. S2b, S2c, S3b**). Western blot analysis revealed that RABIN is strongly enriched in the brain relative to other tissues (**Fig. 1b**). Within the brain, RABIN was predominantly expressed in forebrain structures, with lower levels in the hypothalamus, thalamus, and cerebellum (**Fig. 1c)** and higher levels in the hippocampal CA3 region (**Fig. 1d**). RABIN immunoreactivity colocalized with the marker of excitatory neurons (VGLUT1) but not with markers of inhibitory (GAD2) or peptidergic (PCSK1) neurons (**Figs 1e, S4**). Subcellular fractionation of mouse hippocampus showed that RABIN is enriched in the synaptosome fraction with synaptic glutamate receptor subunits, thereby confirming effective fractionation (**Fig. 1f**). Its presence in synaptoneurosomes, strong colocalization with VGLUT1, and dense neuropil staining suggested presynaptic localization. Consistent with this, dual-immunogold labeling in the hippocampal CA3 region revealed colocalization of RABIN and VGLUT1 in presynaptic terminals **(Fig. 1g)**. Using a SV glycoprotein 2A (SV2A) pull-down assay^25^, we found that RABIN was further enriched in SV fractions with Synaptotagmin 1 (SYT1), RAB3A, and RAB8A **(Fig. 1h)**. Together, these findings place RABIN in the SV pool within glutamatergic neurons.

Based on its AlphaFold-predicted structure (**Fig. 1a**) and similarity to other leucine-rich repeat– containing (LRRC) proteins, we hypothesized that RABIN functions as a noncatalytic scaffold that mediates protein–protein interactions. To define its biochemical role, we first mapped the RABIN protein interactome by using the anti-RABIN antibody. RABIN was immunoprecipitated from wild-type mouse whole-brain preparations, and associated proteins were identified by deep-proteome profiling using liquid chromatography data–independent acquisition mass spectrometry. After normalization to IgG controls, Rab GTPases emerged as the most highly enriched protein class (**Fig. 1i**). Rab binding was validated by reciprocal pulldowns between HA-tagged RABIN and FLAG-tagged RAB3A or RAB8A expressed in primary hippocampal neurons. Because Rab GTPases cycle between the GTP- and GDP-bound states, we tested whether this interaction was nucleotide dependent. Co-immunoprecipitation revealed a strong preference of RABIN for GTP-loaded RAB3A and RAB8A (**Fig. 1J, Fig. S5**). Direct binding in a GTP-dependent manner was further confirmed *in vitro* by using recombinant proteins (**Fig. S6**). Given the conserved role of Rab GTPases in vesicle biogenesis, trafficking, and membrane fusion, these findings suggested that RABIN participates in a previously unrecognized regulatory pathway controlling synaptic transmission. To investigate RABIN’s physiological function, we produced *Rabin^−/−^* mice (**Fig. S3a**). Germline *Rabin* deletion resulted in developmental abnormalities, including reduced brain size and cleft palate (**Fig. S7**), and caused fully penetrant embryonic lethality by embryonic day (E) 18.5. To circumvent lethality, we examined synaptic function in primary hippocampal neurons isolated from E18 embryos. Voltage clamp recordings revealed increased frequency of miniature excitatory postsynaptic currents (mEPSC) but unchanged amplitude in *Rabin^−/−^* neurons compared to controls (**Fig. 1k**). Imaging with the genetically encoded fluorescent glutamate sensor iGluSnFr^26^ showed an increased frequency of spontaneous glutamate release in *Rabin^−/−^*neurons, which was consistent with a presynaptic mechanism, mirrored the electrophysiological findings, and supported a role for RABIN in presynaptic regulation of neurotransmitter release (**Fig. 1l**).

### RABIN constrains quantal glutamate release in excitatory neurons

The perinatal lethality of *Rabin*^−/−^ mice prompted us to generate two conditional alleles enabling Cre-dependent deletion (*Rabin^fl/fl^*) and overexpression (*Rabin^cOE^*) (**Figs. S3b, c**). Using the forebrain-expressed *Emx1^Cre^* driver, we validated Cre-mediated recombination at both alleles by assessing RABIN expression across forebrain regions and in primary cultures of hippocampal neurons (**Figs. S8a, b**). Both *Emx1^Cre^*;*Rabin^fl/fl^* mice and *Emx1^Cre^;Rabin^cOE^* mice were viable and displayed normal growth into adulthood, compared to littermate controls (**Fig. S9**). Consistent with *Rabin^−/−^* neurons in culture, primary neurons from *Emx1^Cre^;Rabin^fl/fl^* mice exhibited an increased frequency of spontaneous glutamate release events at individual synapses, as measured by iGluSnFr (**Figs. 2a, b**). In adult *Emx1^Cre^;Rabin^fl/fl^* brains, variable hippocampal degeneration (discussed below) precluded reliable hippocampal recordings; therefore, electrophysiological analyses were performed in neocortical pyramidal neurons. *Ex vivo* recordings revealed increased mEPSC frequency with unchanged amplitude (**Fig. 2c**). Consistent with presynaptic localization of RABIN to glutamatergic terminals, miniature inhibitory postsynaptic currents (mIPSC) in *Emx1^Cre^;Rabin^fl/fl^*pyramidal neurons were indistinguishable from those in littermate controls (**Fig. 2d**). To assess whether these effects depended on developmental timing or cell-type specificity, we conditionally deleted *Rabin* by using a *Camk2a^Cre^* driver line, which acts later in development and more selectively in forebrain excitatory neurons^27^. *Camk2a^Cre^;Rabin^fl/fl^*mice also showed elevated mEPSC frequency in CA1 pyramidal neurons but normal mEPSC amplitude and mIPSC frequency and amplitude (**Fig. S10**). In contrast, neither optical nor electrophysiological measures of quantal synaptic transmission differed significantly in *Emx1^Cre^;Rabin^cOE^* neurons in culture or CA1 pyramidal neurons, when compared to littermate controls (**Figs. 2e-h**).

**Fig. 2.**
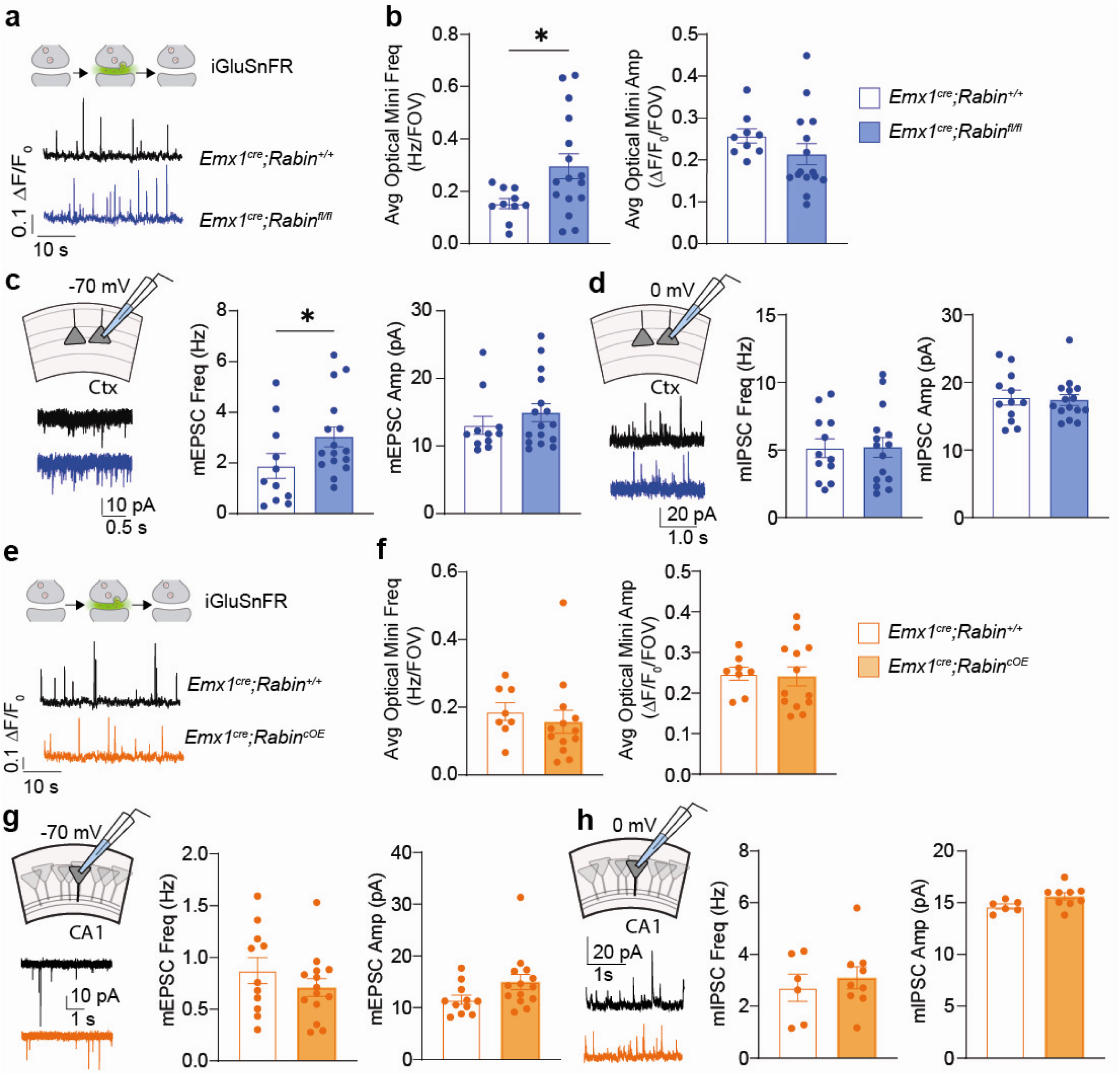
RABIN constrains quantal glutamate release in excitatory neurons. **a**. Representative synaptic iGluSnFr traces from primary hippocampal neurons isolated from *Emx1^Cre^;Rabin^+/+^* mice and *Emx1^Cre^*;*Rabin^fl/fl^* mice. **b**. Optical mini frequency and amplitude (averaged across FOVs); Frequency: Welch’s two-tailed *t*-test, *t*=2.751, \**p*=0.013, n=228, 334 synapses (10, 16 FOVs). Amplitude: Mann-Whitney *U* test, *U*=36, *p*=0.0637, n=193, 329 synapses (9, 15 FOVs). **c.** Representative traces and quantification of miniature excitatory postsynaptic current (mEPSC) frequency and amplitude recorded in *ex vivo* brain slices from the neocortex of *Emx1^Cre^*;*Rabin^+/+^*mice and *Emx1^Cre^*;*Rabin^fl/fl^* mice; Frequency: Mann-Whitney *U* test, *U*=46, \**p*=0.038, n=11, 16 cells. Amplitude: Mann-Whitney *U* test, *U*=69, *p*=0.368, n=11, 16 cells. **d**. Representative traces and mean miniature inhibitory postsynaptic current (mIPSC) frequency and amplitude recorded in *ex vivo* hippocampal CA1 brain slices from *Emx1^Cre^*;*Rabin^+/+^* mice and *Emx1^Cre^*;*Rabin^fl/fl^* mice. Frequency: Mann-Whitney *U* test, *U*=9, *p*=0.1014, n=7, 6. Amplitude: Welch’s two-tailed *t*-test, *t*=0.5922, *p*=0.5661, n=7, 6. **e**. Representative synaptic iGluSnFr traces from primary hippocampal neurons isolated from *Emx1^Cre^*;*Rabin^+/+^* mice and *Emx1^Cre^*;*Rabin^cOE^*mice. **f**. Optical mini frequency and amplitude; Frequency: Welch’s two-tailed *t*-test, *t*=0.708, *p*=0.488, Amplitude: Welch’s two-tailed *t*-test, *t*=0.2291, *p*=0.8213; *n*= 194, 345 synapses (8, 13 FOVs). **g**. Mean mEPSC frequency and amplitude recorded in *ex vivo* hippocampal CA1 brain slices from *Emx1^Cre^*;*Rabin^+/+^*mice and *Emx1^Cre^*;*Rabin^cOE^* mice. Frequency: Mann-Whitney *U* test, *U*=68, *p*=0.648. Amplitude: Welch’s two-tailed *t*-test, *t*=2.00, *p*=0.059. n=11, 14. **h**. Mean mIPSC frequency and amplitude recorded in ex vivo brain slices from *Emx1^Cre^*;*Rabin^+/+^* mice and *Emx1^Cre^;Rabin^cOE^* mice. Frequency: Welch’s two-tailed *t*-test, *t*=0.565, *p*=0.5840, n=6, 9. Amplitude: Welch’s two-tailed *t*-test, *t*=0.839, *p*=0.4133, n=7, 12. Averaged data are presented as the mean ± SEM.

### RABIN suppresses evoked glutamatergic exocytosis by inhibiting Rab GTPase–effector coupling

In their GTP-bound active state, Rab GTPases, including RAB3 and RAB27, regulate active-zone tethering and SV membrane fusion via presynaptic effectors. In particular, vesicle priming and docking require interaction of GTP-loaded RAB3A and its effector RIM1α^11^. To assess the impact of RABIN on this interaction, we purified a recombinant maltose-binding protein fusion of the N-terminal (2-206) RIM1α fragment [MBP–RIM1α (2-206)] that mediates RAB3A binding. Adding recombinant RABIN to GTPγS-loaded RAB3A but not to GDP-loaded RAB3A blocked its interaction with MBP–RIM1α (2-206), indicating that RABIN prevents the binding of RAB3A-GTP to RIM1α (**Fig. 3a**). Several other Rab GTPases, including RAB8A, which regulates the trafficking of trans Golgi-to-plasma membrane, also co-immunoprecipitated with RABIN (**Figs. 1i, j**). Although RAB8A effectors in the brain are largely undefined, we leveraged the established RAB8A–OCRL (539-900) interaction^28^ to test RABIN’s effect^29^. Recombinant RABIN blocked the interaction of GTPγS-loaded RAB8A with GST-tagged OCRL (539-900) (**Fig. 3b**), which was consistent with a broader role for RABIN in inhibiting Rab-effector binding.

**Fig. 3.**
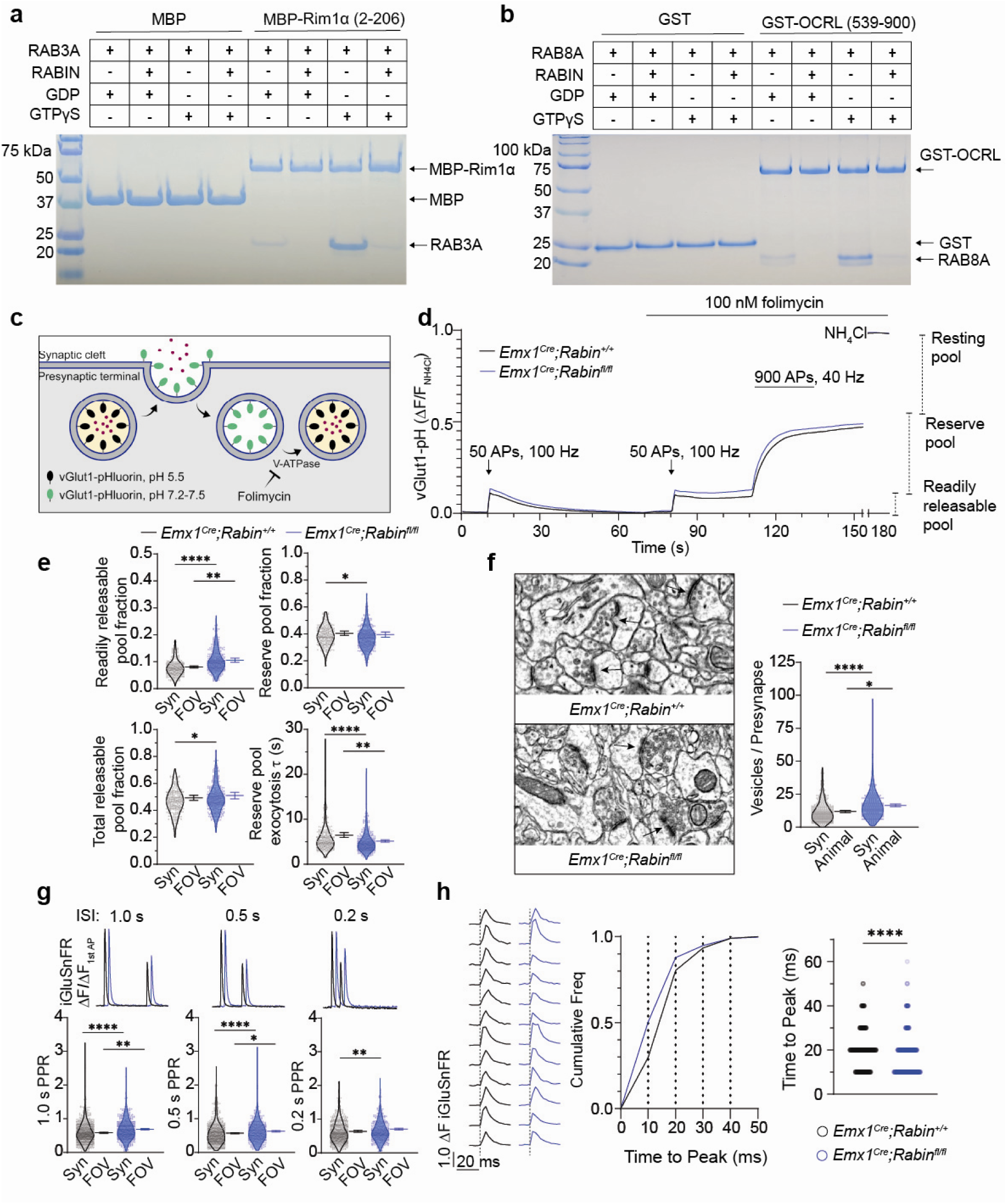
RABIN inhibits RAB GTPase–effector coupling to suppress evoked glutamatergic exocytosis. **a.** *In vitro* pulldown of MBP–RIM1α (2-206) shows that RABIN disrupts GTP-dependent RAB3A–RIM1α binding. **b**. *In vitro* pulldown of GST-OCRL (539-900) displays disruption of GTP-dependent RAB8A-OCRL (539-900) binding by RABIN. **c**. Schematic of the vGlut1-pHluorin sensor measurement of SV exocytosis. **d**. Average of vGlut1-pHluorin fluorescence during stimulation protocol. n=302, 376 synapses. **e**. Quantification of the SV pool parameters extracted from the time courses in (d). Readily releasable pool (RRP) fraction: Mann-Whitney *U* test, *U*=33245, \*\*\*\**p*<0.0001, n= 302, 376 synapses. Welch’s *t*-test*, t*=2.907, \*\**p=*0.0077, n=15,17 FOVs. Reserve pool fraction: Mann-Whitney *U* test, *U*=50935, \**p*=0.0212, n=302, 376 synapses. Welch’s *t*-test*, t*=0.4111, *p*=0.6840, n=15,17 FOVs. Total releasable pool fraction: Mann-Whitney *U* test, *U*=51558, \**p*=0.0396, n=302, 376 synapses. Welch’s *t*-test*, t*=0.5305, *p*=0.5998, n=15,17 FOVs. Reserve pool exocytosis rate: Mann-Whitney *U* test, *U*=44106, \*\*\*\**p*<0.0001, n=302, 376 synapses. Mann-Whitney *U* test*, U*=56.5, \**p*=0.0063, n=15,17 FOVs. **f**. (Left) Electron micrographs of hippocampal CA1 synapses and associated SVs. (Right) Quantification of the SVs within each synapse for the indicated genotypes. Mann-Whitney *U* test, *U*=28772, \*\*\*\**p* <0.0001, n=333, 255 vesicles; Welch’s *t*-test*, t*=3.468, \**p=*0.0256, n=3, 3 animals. **g**. (Top) representative iGluSnFr fluorescence responses to two field stimuli at 1.0-, 0.5-, or 0.2-s inter-stimulus intervals (ISIs). The traces for each genotype are offset 0.1 s for clarity. (Bottom) The ratios of peak iGluSnFr fluorescence of the second response were normalized to the initial response and plotted as the paired-pulse ratio (PPR) for each synapse at the indicated ISI. The 1.0-s ISI: Mann-Whitney *U* test, *U*=173021, \*\*\*\**p*<0.0001, n=786, 536 synapses. Welch’s two-tailed *t*-test, *t*=3.585, \*\**p=*0.0016, n=17,13 FOVs. The 0.5-s ISI: Mann-Whitney *U* test, *U*=131807, \*\*\*\**p*<0.0001, n=729,420 synapses. Welch’s two-tailed *t*-test, *t*=2.079, \**p=*0.0480, n=17,13 FOVs. The 0.2-s ISI: Mann-Whitney *U* test, *U*=128496, \*\**p*<0.0088, n=682,416 synapses. Welch’s two-tailed *t*-test, *t*=1.718, *p=*0.0969, n=17,13 FOVs. **h**. (Left) representative iGluSnFr fluorescence traces measured at single synapses immediately after a field stimulus (indicated by the dashed vertical lines) for each indicated genotype. (Middle) Cumulative distribution plot of the time-to-peak fluorescence for all synapses. (Right) time-to-peak fluorescence is plotted for each synapse and compared between the indicated genotypes. Mann-Whitney *U* test, *U*=208809, \*\*\*\**p*<0.0001, *n* = 641, 831 synapses.

RIM1α, a core active-zone constituent, mediates SV docking, priming, and recruitment of voltage-gated calcium channels^1,30^. Given RABIN’s inhibition of active RAB3A–RIM1α (2-206) binding, we assessed its role in SV exocytosis. We expressed the fluorescent pH-sensitive reporter of glutamatergic SV turnover vGlut1-pHluorin^31^ in primary hippocampal neurons from *Emx1^Cre^;Rabin^fl/fl^* mice and littermate controls (**Fig. 3c**). Both the size of the readily releasable pool (RRP) and the rate of the reserve pool exocytosis were increased in *Emx1^Cre^;Rabin^fl/fl^* synapses compared with controls (**Figs. 3d, e**). To determine whether the increased release rate was a general effect, rather than frequency-specific, we measured SV turnover in synapses from rest to a range of stimulation frequencies. Experiments were performed in the absence of the specific inhibitor of vacuolar-type H^+^-ATPase folimycin to track frequency-dependent endocytosis. *Emx1^Cre^;Rabin^fl/fl^*neurons exhibited an increased exocytosis rate that saturated above 20 Hz, and endocytosis was unchanged (**Fig. S11**).

Electron microscopy (EM) in the CA1 region revealed more SVs in *Emx1^Cre^;Rabin^fl/fl^*synapses than in those of littermate controls, whereas synapse number, size, active-zone vesicle number, and vesicle pool density were unchanged; vesicle density was reduced in *Emx1^Cre^;Rabin^cOE^*mice (**Fig. 3f, Table S2**). These data indicate that RABIN constrains SV pool size and the RRP without affecting the number of synapses.

Increased vesicle numbers could affect presynaptic short-term plasticity by changing the rate of vesicle replenishment at the active zone. Field-evoked paired pulse depression (PPD), which is measured with iGluSnFr, showed greater recovery in *Emx1^Cre^;Rabin^fl/fl^*synapses and the opposite phenotype in *Emx1^Cre^;Rabin^OE^*synapses (**Figs. 3g, S12**). Time-to-peak analysis of iGluSnFr signals revealed faster release kinetics in *Emx1^Cre^;Rabin^fl/fl^* synapses and slower kinetics in *Emx1^Cre^;Rabin^cOE^* synapses (**Figs. 3h, S13**). Together, these data indicate that RABIN acts as a presynaptic negative regulator of Rab GTPase-mediated signaling and SV turnover. Modulating *Rabin* gene dosage bidirectionally altered the spontaneous quantal and evoked synaptic transmission. These convergent effects are consistent with dysregulated Rab–effector coupling at the presynaptic active zone, thereby prompting our investigation of the behavioral and pathological consequences of Rab–GTPase dysregulation caused by *Rabin* deficiency.

### Forebrain *Rabin* deficiency confers epileptiform pathology

Histological analysis revealed hippocampal degeneration in ∼30% of *Emx1^Cre^;Rabin^fl/fl^*mice, resembling sclerosis associated with temporal lobe epilepsy^32^ (**Fig. S14**). Degeneration was most prominent throughout the dorsal and ventral CA3 region, extending into CA1 while largely sparing CA2, except in the most extreme cases (**Fig. S14a**). In a subset of animals, necrosis spread across the CA3–CA1 regions and into the neocortex; this was accompanied by foamy macrophages, a hallmark of neurodegeneration and demyelination (**Figs. S14b, c**). We hypothesized that this tissue damage arises from excitotoxicity driven by enhanced glutamate release in *Emx1^Cre^;Rabin^fl/fl^* mice. Therefore, even in the absence of overt sclerosis, we observed chronic neuroinflammation characterized by increased astrocyte and microglia density in the hippocampus and neocortex (**Figs. S15a, b**). Consistent with early neurodegenerative processes, the expression of cleaved caspase-3, a marker of apoptotic signaling, was also elevated (**Fig. S15c**). Brains exhibiting extensive necrosis with dense GFAP, IBA1, and cleaved caspase-3 labeling were excluded from quantitative analysis. In contrast, *Camk2a^Cre^;Rabin^fl/fl^*mice, in which *Rabin* was deleted later in development, showed no overt hippocampal degeneration (**Fig. S16**).

We hypothesized that excessive glutamate release in RABIN-deficient mice leads to seizures. Therefore, we assessed *Emx1^Cre^;Rabin^fl/fl^*mice and littermate controls by using behavioral assays associated with epileptic phenotypes. We identified increased locomotor activity in the open field, heightened risk taking in the elevated plus maze, and reduced social interaction in the social preference test (**Fig. S17a-c**). *Camk2a^Cre^;Rabin^fl/fl^*mice showed similar deficits in the open field and social preference assays (**Fig. S18a-c**). These behavioral deficits closely resembled those reported in established rodent models of epilepsy^33,34^. Based on these behavioral and histopathological phenotypes, and the enhanced glutamatergic synaptic transmission, we hypothesized that *Rabin* loss confers epileptic susceptibility. Consistent with this, *Emx1^Cre^;Rabin^fl/fl^*mice exhibited a reduced threshold for kainic acid (KA)-induced seizures at doses (15-20 mg/kg) that triggered status epilepticus in fewer than 50% of controls (**Fig. 4a**). To characterize seizure activity electrophysiologically, we performed long-term video-EEG telemetry in freely behaving mice. At a subthreshold KA dose (10 mg/kg), *Emx1^Cre^;Rabin^fl/fl^* mice had greater seizure susceptibility, which was quantified as a larger increase in EEG power across frequency bands (**Fig. 4b, c**). We observed spontaneous epileptiform discharges and seizures in *Emx1^Cre^;Rabin^fl/fl^*mice before KA administration that persisted after washout, but both were absent in littermate controls (**Figs. 4d, e; Movie S1**). Similarly, we observed greater seizure susceptibility when *Rabin* was deleted later in development in *Camk2a^Cre^;Rabin^fl/fl^* mice. *Camk2a^Cre^;Rabin^fl/fl^*mice exhibited a reduced threshold for KA-induced seizures at subthreshold doses (**Fig. S19a**) and a larger increase in EEG power across frequency bands (**Fig. S19b**). We also observed spontaneous epileptiform discharges and seizures in *Camk2a^Cre^;Rabin^fl/fl^*mice before KA administration that persisted after washout, but both were absent in littermate controls (**Fig. S19d, e**).

**Fig. 4.**
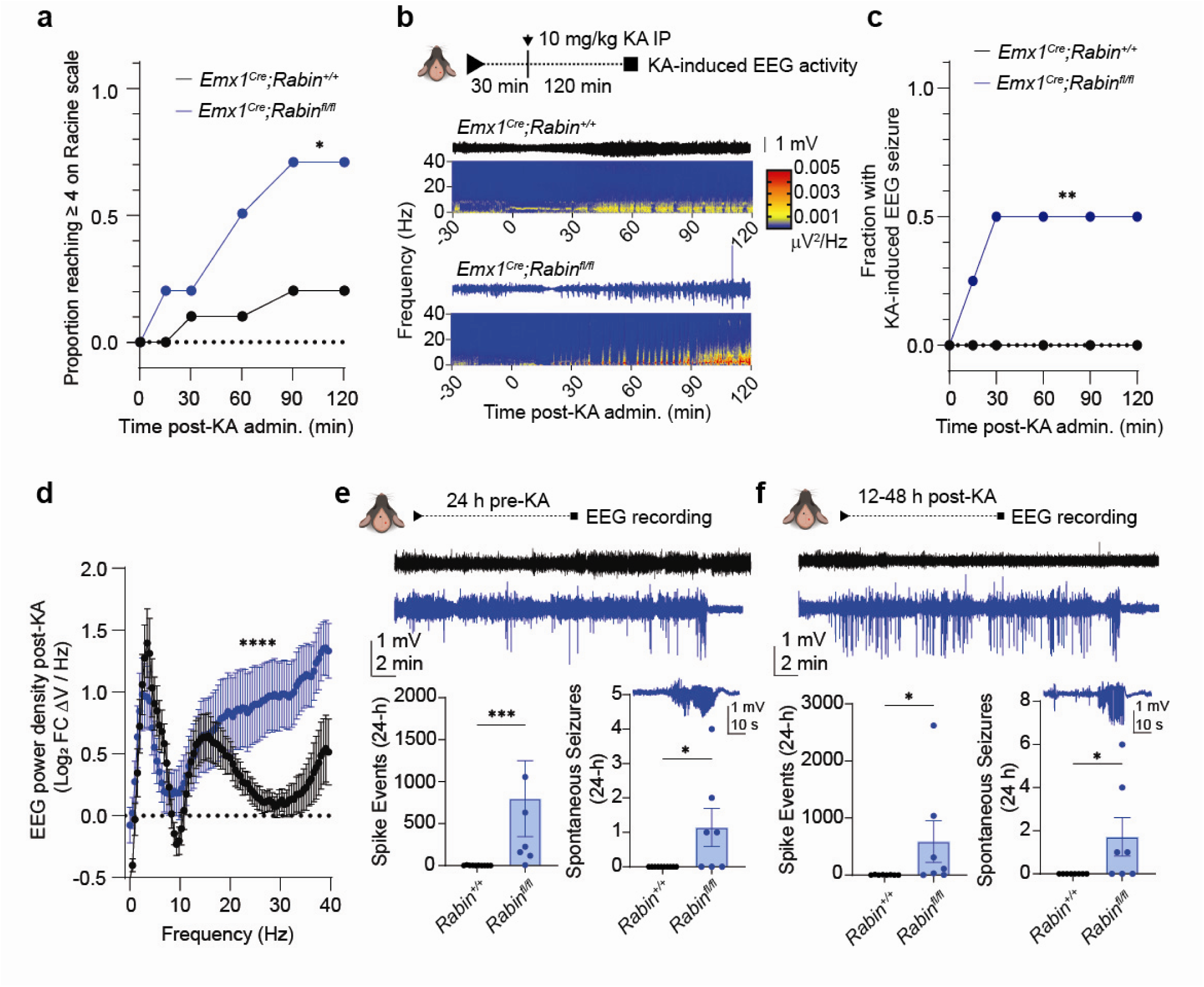
*Rabin* deficiency in the forebrain confers epileptiform pathology. **a.** *Rabin* deletion lowers the threshold for behavioral seizures. *Emx1^Cre^*;*Rabin^fl/fl^*(blue plot) and littermate controls (black plot) were injected intraperitoneally with a subthreshold dose of kainic acid (KA). Seizure severity was scored at 30-min intervals using the Racine scale, and the proportions of total animals reaching stage ≥4 seizures were quantified. Two-way ANOVA, *F*(1, 5)=10.55, \**p*=0.0228, n=10, 10 mice. **b**. Electroencephalography (EEG) telemetry showed increased EEG power in *Emx1^Cre^*;*Rabin^fl/fl^* mice relative to controls after subthreshold KA administration. Representative EEG traces and power spectra are shown for control and *Emx1^Cre^*;*Rabin^fl/fl^*mice for the entire period from 30 min before to 120 min after injection. **c**. *Emx1^Cre^*;*Rabin^fl/fl^* exhibit EEG seizure activity in response to a subthreshold dose of KA, whereas littermate controls displayed no EEG abnormalities. Two-way ANOVA revealed a significant effect of genotype *F*(1,5) =19.29, \*\**p*=0.0071; *n*=10, 8 mice. **d**. *Emx1^Cre^*;*Rabin^fl/fl^*mice show a greater increase in EEG power across the full frequency spectrum measured. Data are plotted as the mean Log_2_ ± SEM. Two-way ANOVA revealed a significant effect of genotype *F*(1,1230)=257.9, \*\*\*\**p*<0.0001, *n*=9, 8 mice. **e**. *Emx1^Cre^*;*Rabin^fl/fl^*mice show spontaneous neuronal discharges and seizures in a 24-h period before KA administration. Representative EEG traces are shown for control (black) and *Emx1^Cre^*;*Rabin^fl/fl^* (blue) mice. The number of spike events and spontaneous seizures are plotted as mean ± SEM. The representative trace above the seizure summary data depicts a spontaneous seizure event in an *Emx1^Cre^*;*Rabin^fl/fl^* mouse. Spike events: Mann-Whitney *U* test, *U*=1, \*\*\**p*=0.0002, n=10, 7. Seizure events: Unpaired one-tailed *t*-test, *t*=2.505, \**p*=0.0121, n=10, 7 mice. **f**. *Emx1^Cre^*;*Rabin^fl/fl^*mice show spontaneous neuronal discharges and seizures after KA washout. Representative EEG traces are shown for control mice and *Emx1^Cre^*;*Rabin^fl/fl^*mice. Data are plotted as in **4e**. The representative trace above the seizure summary data depicts a seizure event in an *Emx1^Cre^*;*Rabin^fl/fl^*mouse. Spike events: Mann-Whitney *U* test, *U*=9.5, \**p*=0.0152, n=9, 7 mice. Seizure events: Unpaired one-tailed *t*-test, *t*=2.201, \**p*=0.0225, n=9, 7 mice. Averaged data are presented as the mean ± SEM.

### Epileptiform pathology is rescued by levetiracetam and *Rabin* overexpression

Epileptiform activity in *Rabin*-deficient mice supports a presynaptic mechanism involving excessive Rab3-dependent vesicle delivery at excitatory synapses. Levetiracetam (LEV), a first-line anti-epileptic drug that targets voltage gated calcium channels and SV2A, was used to test this model^35–37^. In cultured neurons, LEV produced a reversible, dose-dependent reduction in vGlut1-pHluorin responses to 100 stimuli at 20 Hz (**Fig. 5a**) and decreased iGluSnFR responses to single stimuli (**Fig. 5b**), consistent with reduced presynaptic release. In vivo, *Emx1^Cre^;Rabin^fl/fl^*mice exhibited spontaneous epileptiform discharges that were significantly reduced by 200 mg/kg LEV (**Fig. 5c**), consistent with a presynaptic mechanism for *Rabin*-dependent pathology.

**Fig. 5.**
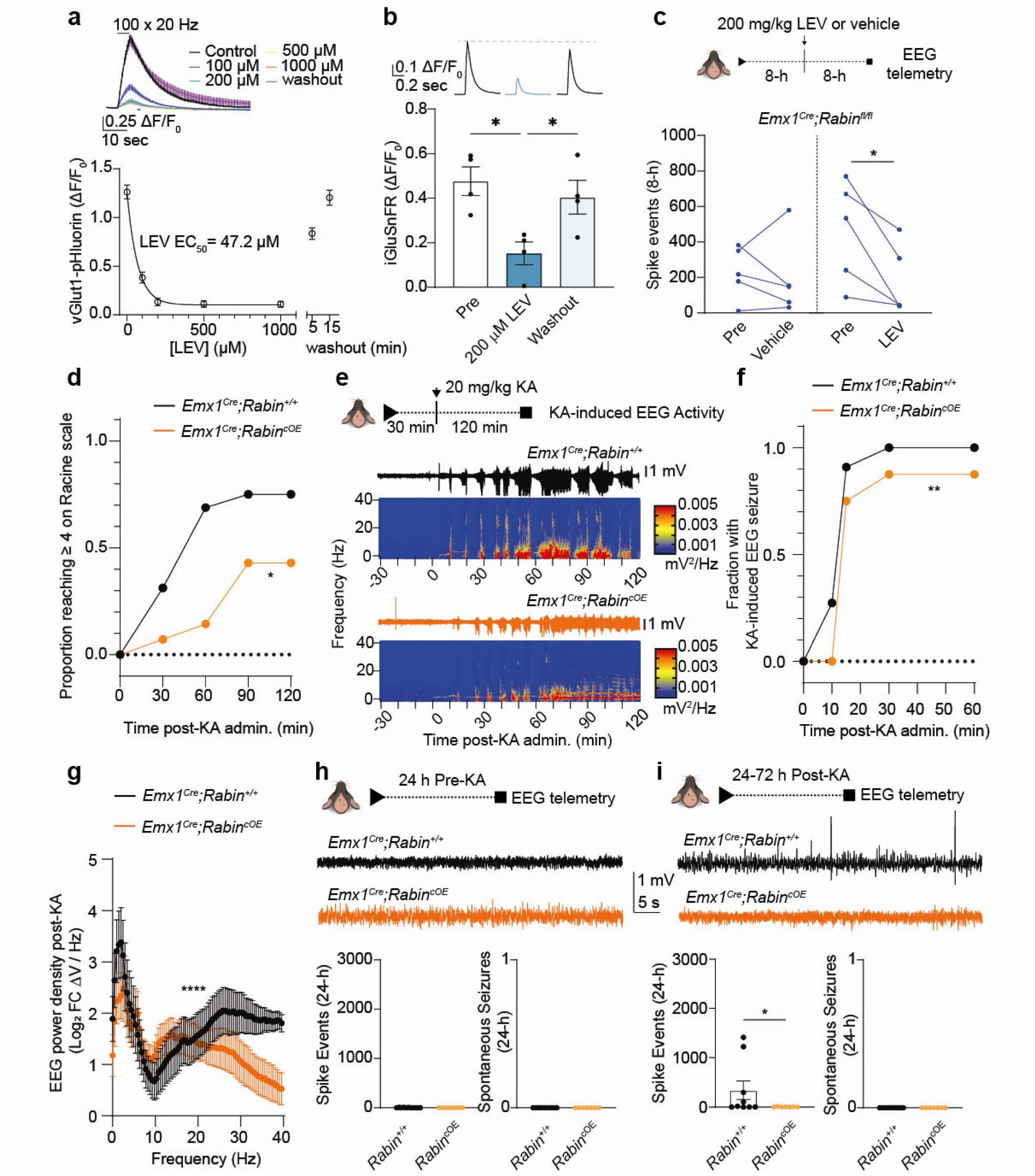
Epileptiform activity is rescued by levetiracetam and Rabin overexpression. **a.** Levetiracetam reversibly and dose-dependently reduces vGlut1-pHluorin responses evoked by 100 stimuli at 20 Hz. n=20 synapses. **b.** Levetiracetam reversibly reduces iGluSnFR responses evoked by field stimulation. Repeated measures one-way ANOVA, pre-LEV: \**p*=0.0163; pre-Washout: *p*=0.2683; LEV-Washout: \**p*=0.0292. n=4 FOVs. **c.** EEG telemetry in *Emx1^Cre^*;*Rabin^fl/fl^*mice shows that levetiracetam significantly reduces spike events during the 8 h following 200 mg/kg i.p. administration. **d**. *Rabin* overexpression increases the threshold for behavioral seizures. *Emx1^Cre^*;*Rabin^cOE^*mice (orange plot) and littermate controls (black plot) received a threshold dose of KA. Seizure severity was scored at 30 min intervals by using the Racine scale, and the proportions of total animals reaching stage ≥4 seizures were quantified. Two-way ANOVA revealed a significant effect of genotype *F*(1,3)=40.11, \*\**p* =0.0080, n=16, 14 mice. **e**. EEG telemetry measured an attenuated increase in EEG power after administration of the threshold dose of KA in *Emx1^Cre^*;*Rabin^cOE^*mice, compared with littermate controls. Representative EEG traces and power spectra are shown for control mice and *Emx1^Cre^*;*Rabin^cOE^*mice during the period from 30 min before to 120 min after KA injection. **f**. EEG seizure activity was reduced in *Emx1^Cre^*;*Rabin^cOE^*mice after KA administration compared to littermate controls. Two-way ANOVA revealed a significant effect of genotype *F*(1,6)=19.51, \*\**p*=0.0045; n=9, 7 mice. **g**. *Emx1^Cre^*;*Rabin^cOE^* mice show an attenuated increase in EEG power across the full frequency spectrum measured. Two-way ANOVA revealed a significant effect of genotype *F*(1,1312)=39.73, \*\*\*\**p* <0.0001, n=10, 8 mice. **h**. *Emx1^Cre^*;*Rabin^cOE^*mice and littermate controls show no epileptic activity in the 24-h period before KA administration. Representative EEG traces are shown for control mice and *Emx1^Cre^*;*Rabin^cOE^*mice. Spike and seizure counts were unchanged in *Emx1^Cre^*;*Rabin^cOE^*mice, Welch’s *t*-test, *t*=1.982, *p*=0.0709, n=11, 8 mice. **i.** Littermate control mice exhibit epileptiform discharges in the 24- to 72-h period after KA washout, but *Emx1^Cre^*;*Rabin^cOE^*mice show no epileptic activity during this period. Representative EEG traces are shown for control mice and *Emx1^Cre^*;*Rabin^cOE^*mice. Data are plotted as in **5d**. Mann-Whitney *U* test, *U*=13, \**p*=0.0497, n=9, 7 mice. Averaged data are presented as the mean ± SEM.

As described above, conditional overexpression of *Rabin* produced synaptic phenotypes that were largely the opposite of those observed after its knockdown, including slowed field-evoked iGluSnFr kinetics and enhanced PPD. Overall, synaptic alterations associated with conditional overexpression of *Rabin* were modest, suggesting that endogenous *Rabin* expression levels are sufficient to saturate its negative modulation of Rab GTPases under basal conditions. Therefore, we hypothesized that increasing *Rabin* expression is beneficial in pathologic states that overwhelm the normal homeostatic control of SV turnover. Accordingly, *Emx1^Cre^;Rabin^cOE^*mice exhibited an increased threshold for KA-induced status epilepticus when challenged with doses that elicited seizures in 100% of the control animals (i.e., 20-25 mg/kg) (**Fig. 5d**). Consistent with this behavioral protection, *Emx1^Cre^;Rabin^cOE^*mice also showed a blunted increase in EEG power after administration of 20 mg/kg KA in females or 25 mg/kg KA in males (**Figs. 5e-g**). As expected, epileptiform activity was not detected in *Emx1^Cre^;Rabin^cOE^*mice during the 24-h period preceding KA injections (**Fig. 5h**). However, *Rabin* overexpression prevented the emergence of prolonged epileptiform discharges during the 24- to 72-h period after KA washout, a pathologic phenotype observed only in control littermates (**Fig. 5i**).

## DISCUSSION

Here we identified LRRC57/RABIN as an SV protein that is enriched in glutamatergic synapses and restrains Rab–effector coupling by blocking effector engagement. Rather than regulating Rab nucleotide state or membrane association, RABIN binds GTP-loaded synaptic Rabs and functions as a noncatalytic decoy that limits effector access. Consequently, deletion of *Rabin* drives excessive glutamate release and epileptiform pathology, and increased *Rabin* expression is protective in a seizure model. Together, these findings define a previously unrecognized mode of presynaptic Rab regulation that preserves excitatory circuit stability.

Rab GTPases act as molecular switches that engage specific effector proteins to control vesicle trafficking, docking, and fusion. Canonical Rab regulation centers on GEFs, GAPs, and GDIs that govern Rab nucleotide state and membrane cycling. This multilayered regulatory network—comprising more than 40 GEFs, more than 40 GAPs, and two GDIs in humans—underscores the importance of regulating the activity of Rab GTPases in virtually all aspects of vesicular biology in eukaryotes, but whether additional mechanisms exist to restrain Rab–effector coupling has remained unclear. Here we describe the founding member of the fifth class of regulators found at presynaptic terminals; an inhibitor of Rab/GTP–effector interaction, which we accordingly named RABIN. RABIN selectively associates with GTP-loaded synaptic Rabs, including RAB3A and RAB8A, and competitively inhibits their interaction with effectors, such as RIM1α and OCRL. To our best knowledge, this is the first example of a decoy-effector mechanism that provides a direct means of limiting Rab output without altering Rab’s activation state, enabling rapid, local control of vesicle mobilization under the conditions of high synaptic demand.

Inhibition of RAB3A–RIM1α coupling by RABIN provides a mechanistic explanation for the presynaptic phenotypes observed after *Rabin* deletion. RIM1α is a well-recognized central organizer of the active zone, which promotes SV docking, priming, and calcium channel recruitment^1,11,30^; RAB3-dependent RIM1α engagement would be expected to shift the balance from vesicle retention at the active zone to mobilization from reserve pools^14,38^. Consistent with this, the loss of RABIN increased the size of the RRP, accelerated vesicle replenishment, and reduced PPD of glutamate release, indicating enhanced availability of release-competent vesicles. Beyond RAB3A, RABIN binds additional synaptic Rabs, suggesting that it functions as a multi-Rab inhibitor within the SV-trafficking pathway.

Our data further revealed a gene-dosage dependence of RABIN function that is consistent with a homeostatic role in synaptic transmission. Deletion of *Rabin* robustly enhanced glutamate release, whereas overexpression produced comparatively modest baseline effects, despite bidirectional changes in short-term plasticity and release kinetics. This asymmetry suggests that endogenous RABIN levels are normally sufficient to saturate Rab GTPase–effector inhibition under physiological conditions, positioning RABIN as a stabilizing constraint that limits excessive vesicle mobilization, rather than as a primary determinant of basal release probability. Pathological phenotypes emerge when synaptic demand exceeds this buffering capacity^2^, as occurs after *Rabin* deletion or during sustained network hyperactivity; increasing RABIN expression preserved synaptic and circuit stability without broadly suppressing neurotransmission.

At the circuit level, *Emx1^Cre^*-mediated loss of RABIN in forebrain excitatory neurons produced epileptiform activity and spontaneous seizures; under conditions of higher penetrance, it caused hippocampal degeneration with prominent neuroinflammation and apoptotic signaling. We also observed epileptiform discharges and seizure susceptibility after *Camk2a^Cre^*-mediated deletion of *Rabin* in excitatory neurons, in the absence of hippocampal damage. This result indicated that disrupting presynaptic Rab regulation within excitatory circuits is sufficient to drive epileptiform activity, and structural pathology likely emerged as a downstream severity-dependent consequence of hyperexcitability, rather than its primary cause. The CA3-predominant hippocampal degeneration observed in a subset of animals closely resembled sclerosis associated with temporal lobe epilepsy^32^, thereby supporting a model in which chronic synaptic release culminates in excitotoxic injury.

Neither activating nor inactivating mutations in the *RABIN* gene have ever been observed in the clinic. This is not surprising, considering the very high degree of evolutionary conservation of Rabin and the fact that a knockout of this gene in mice and invertebrate models results in lethality. In contrast, inactivating mutations in synapsins I and II, which are well-known regulators of vesicular transport, have been observed in many patients with epilepsy^39–42^. Deletion of Rab3a robustly rescues the epileptic phenotypes observed in *Syn II*-knockout mice^43,44^. Therefore, although *RABIN* does not function as an etiologic disease gene, our results strongly support the idea that it can act as an epistatic modifier whose modulation may be a crucial for controlling synaptic stability and have therapeutic potential— analogous to PCSK9 inhibition in the treatment of hypercholesterolemia in atherosclerotic cardiovascular disease^45,46^. Hence, the “molecular glue” type of small molecules^47,48^ for orthosteric stabilization of the RABIN–RAB3 interaction would have a beneficial therapeutic effect in patients with epilepsy, and small molecules that prevent RABIN–RAB·GTPase interaction would be beneficial to enhance glutamate release in psychiatric diseases in which glutamatergic signaling is impaired, such as major depressive disorders^49^ or schizophrenia^50,51^.

Together, these findings position RABIN as a new class of presynaptic Rab regulator that constrains SV turnover to preserve circuit balance and integrity. More broadly, they suggest that bidirectional fine-tuning of Rab–effector coupling, rather than globally altering neurotransmitter release, represents an effective strategy for stabilizing hyper- or hypoexcitable neural circuits in various pathologic states. Future work will be required to define how RABIN is regulated by neural activity to map its full complement of synaptic Rab targets, including whether RABIN constrains Rab signaling outside the presynaptic compartment. Whether similar decoy-effector mechanisms operate at other trafficking steps or in distinct neuronal populations also remains to be determined.

## Acknowledgements

Research reported in this publication was supported by the National Institute of Mental Health of the National Institutes of Health under Award Number R21MH138869. This work was also supported by the American Lebanese Syrian Associated Charities (ALSAC). The St. Jude Center for Advanced Genome Engineering, Genetically Engineered Mouse Models Shared Resource, and protein production facility were funded by the National Cancer Institute (P30CA021765). We thank the Zakharenko lab members for constructive comments; Dr. Valerie Stewart for help with mutant mouse production; Maria Gulinello and Kerry Heath for assistance with mouse behavioral assays; George Campbell for training and microscope assistance; Meifen Lu for assistance with immunohistochemistry and immunofluorescence assay development; Cai Li for assistance with statistical analysis; Dr. Ti-Cheng Chang for assistance with bioinformatic analysis; and Dr. Angela McArthur for manuscript editing. The content is solely the responsibility of the authors and does not necessarily represent the official views of the National Institutes of Health or other granting agencies.

## Author Contributions

Conceptualization, D.G., J.G.M., S.S.Z.; investigation, D.G., J.G.M., B.J.W.T., C.M.D., C.D.S., L.J.J., W.J.C, J. K., Z.W., S.P., M.M., J.Y.F., D.A.L., T.C.; methodology, M.J., C.G.R., R.J.H., J.D.V., J.P., S.M.P.-M.; resources, funding, and supervision, S.S.Z.; writing—original draft preparation, J.G.M., D.G.; writing—review and editing, S.S.Z, J.G.M, D.G. All authors have read and agreed to the published version of the manuscript.

## Data and materials availability

Data are available upon request from the corresponding author. Proteomics data are available at PRIDE (Proteomics IDEntification Database), dataset identifier PXD072971 (username; password: zIK1EXQX12B0).

## SUPPLEMENTARY INFORMATION

**Supplementary Figure S1.**
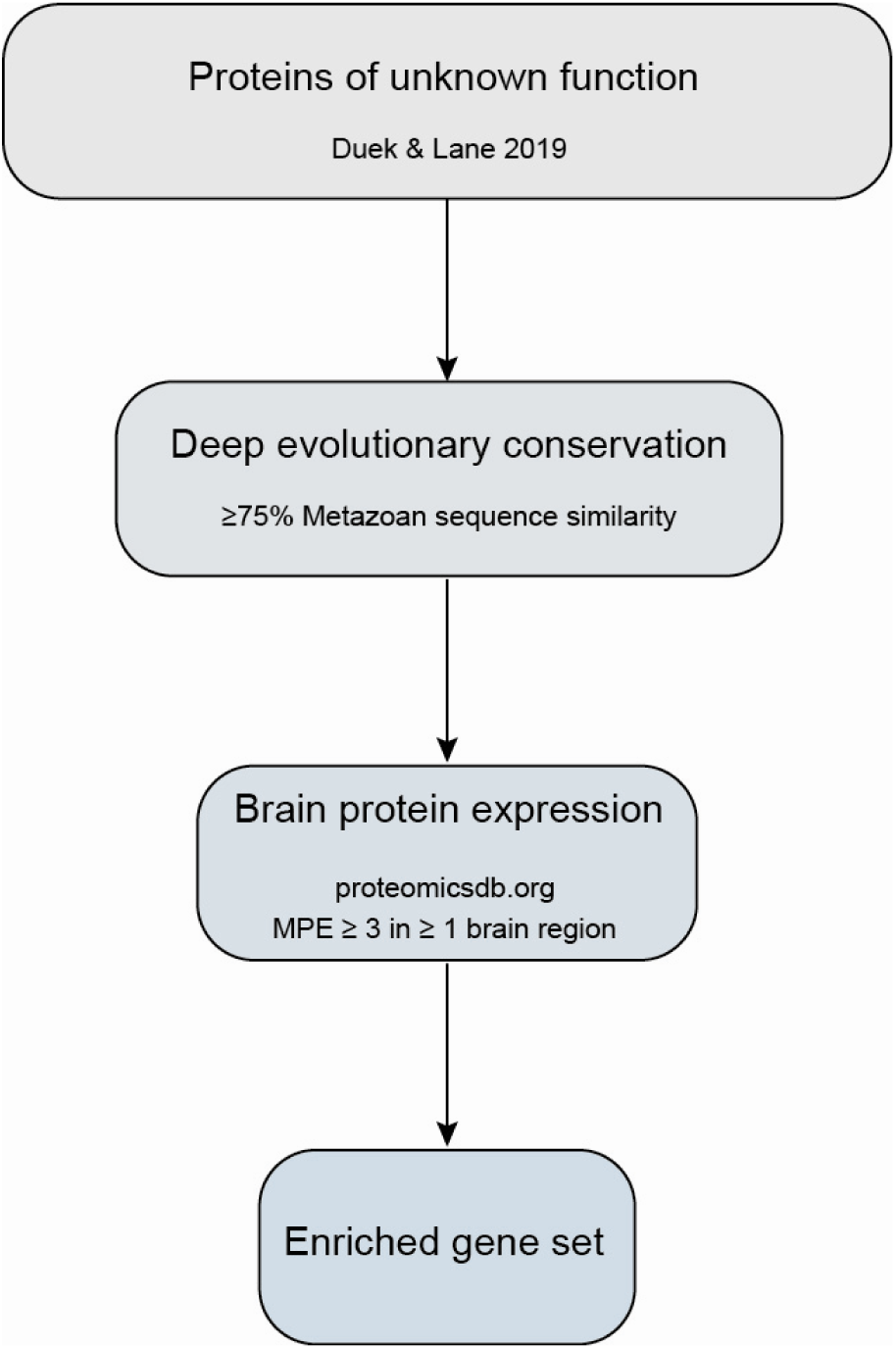
An enrichment-based bioinformatic pipeline for identifying brain-enriched genes. MPE, Median Protein Expression.

**Supplementary Figure S2.**
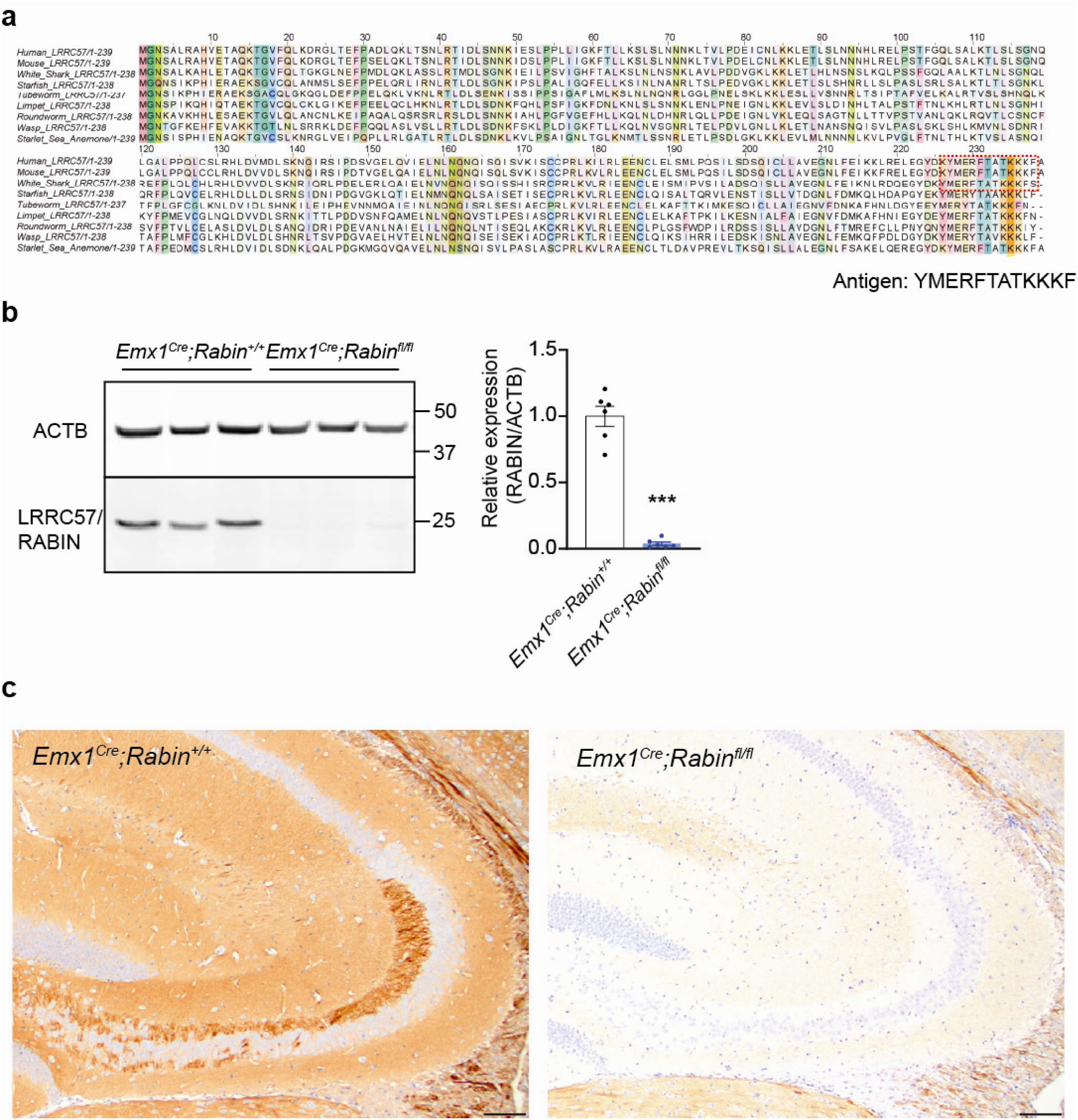
Development and validation of an LRRC57/RABIN monoclonal antibody. **a.** Evolutionary conservation of the LRRC57/RABIN protein across species. **b.** Immunoblot validation of *Lrrc57* deletion in hippocampal lysates from *Emx1^Cre^;Rabin^fl/fl^*mice (blue plot), relative to that in littermate controls (black plot). Welch’s two-tailed *t*-test, *t*=12.57, \*\*\**p* <0.0001, n=6, 6 mice. **c.** Immunohistochemical validation of LRRC57/RABIN staining in the CA3 region of the hippocampus from a control mouse and an *Emx^Cre^;Rabin^fl/fl^* littermate. Scale bars, 100 μm. Averaged data are presented as the mean ± SEM.

**Supplementary Figure S3.**
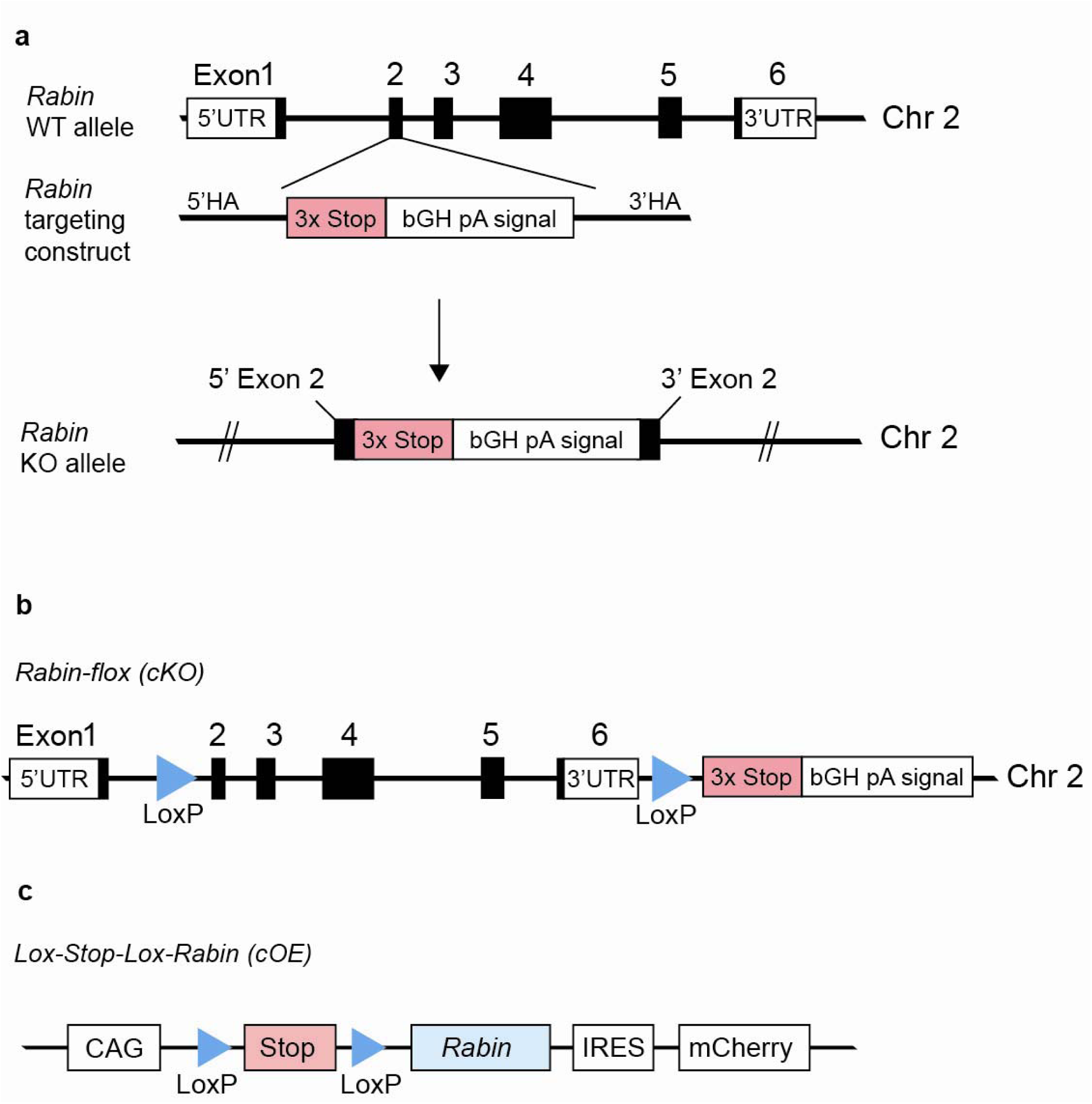
Allele design for germline and conditional knockout (cKO) and conditional overexpression (cOE) mouse lines. **a.** Germline knockout mice were developed by disrupting exon 2 with 3 stop codons, followed by a bovine growth hormone polyadenylation (bGH pA) signal to avoid engagement of the nonsense-mediated decay pathway for resulting truncated mRNA. **b.** A CRE-dependent conditional *Rabin* allele was developed by inserting two *LoxP* sites (blue triangles) flanking exon 2 and the 3’UTR of the *Rabin* gene, followed by 3 stop codons and a polyadenylation signal. **c**. Genomic organization of the *Rabin* conditional overexpression transgene. In the presence of Cre recombinase, the CAG promoter drives the bicistronic expression of *Rabin* and mCherry fluorescent protein.

**Supplementary Figure S4.**
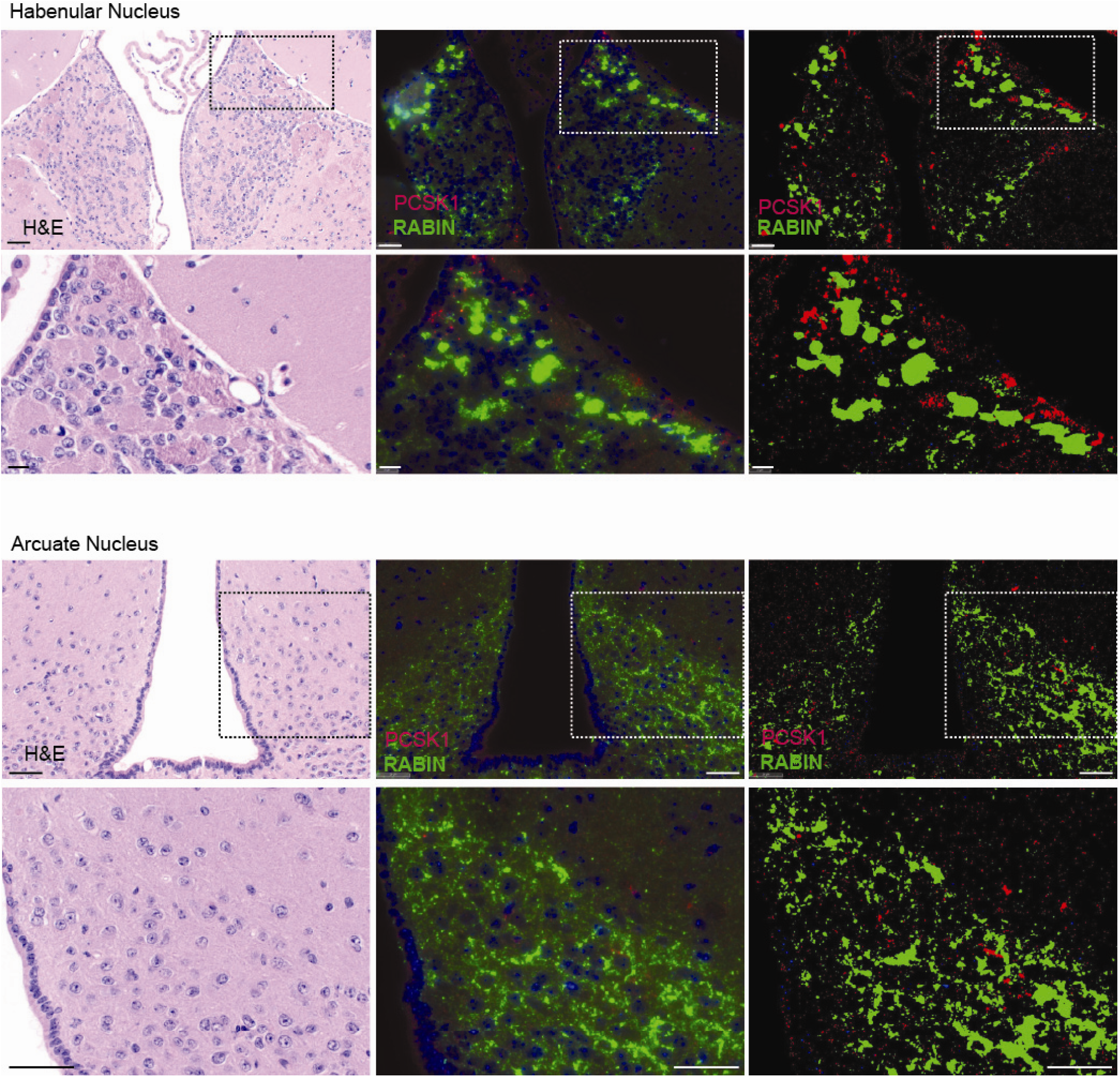
RABIN protein is not expressed in peptidergic neurons. (Left) H&E staining shows the anatomy and location of immunostaining in the habenular and arcuate nuclei. (Middle) Co-immunolabeling of RABIN (green) and PCSK1 (red), a prohormone convertase and marker of peptidergic neurons. (Right) Image contrast enhancement and segmentation shows minimal overlap between RABIN and PCSK1. Scale bars, 100 μm, inset, 50 μm.

**Supplementary Figure S5.**
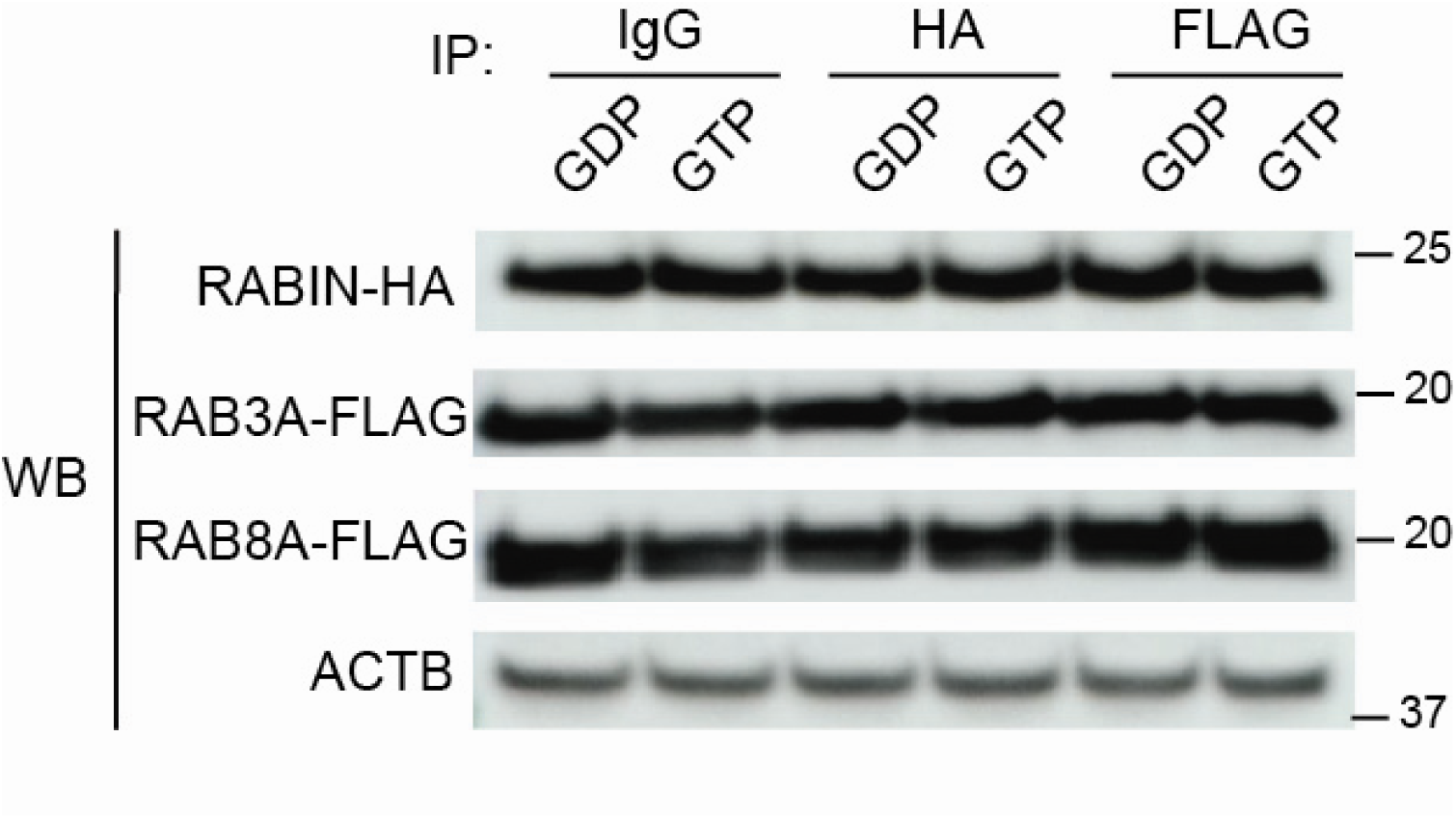
Loading controls for the Western blot (WB) pull-down experiment in. **Fig. 1j**. Extracts were prepared from primary hippocampal neurons transduced with *Rabin-HA* and either *Rab3a-FLAG* or *Rab8a-FLAG*. Representative band densities are shown for RABIN-HA, RAB3A-FLAG, RAB8A-FLAG, and endogenous β-actin (ACTB) expression.

**Supplementary Figure S6.**
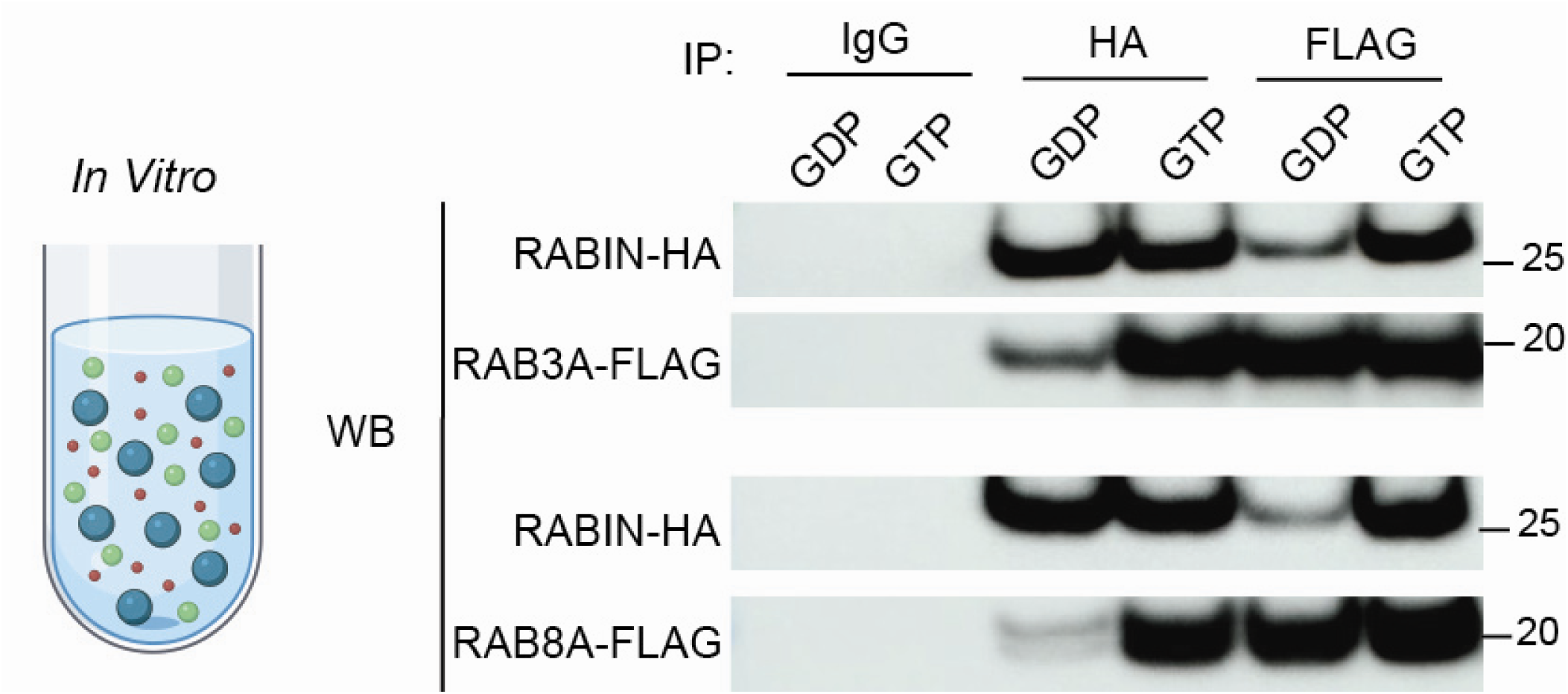
GTP dependence of direct binding between recombinant RAB3A-FLAG and RAB8A-FLAG with HA-RABIN is shown by co-elution *in vitro* after reciprocal HA or FLAG immunoprecipitation.

**Supplementary Figure S7.**
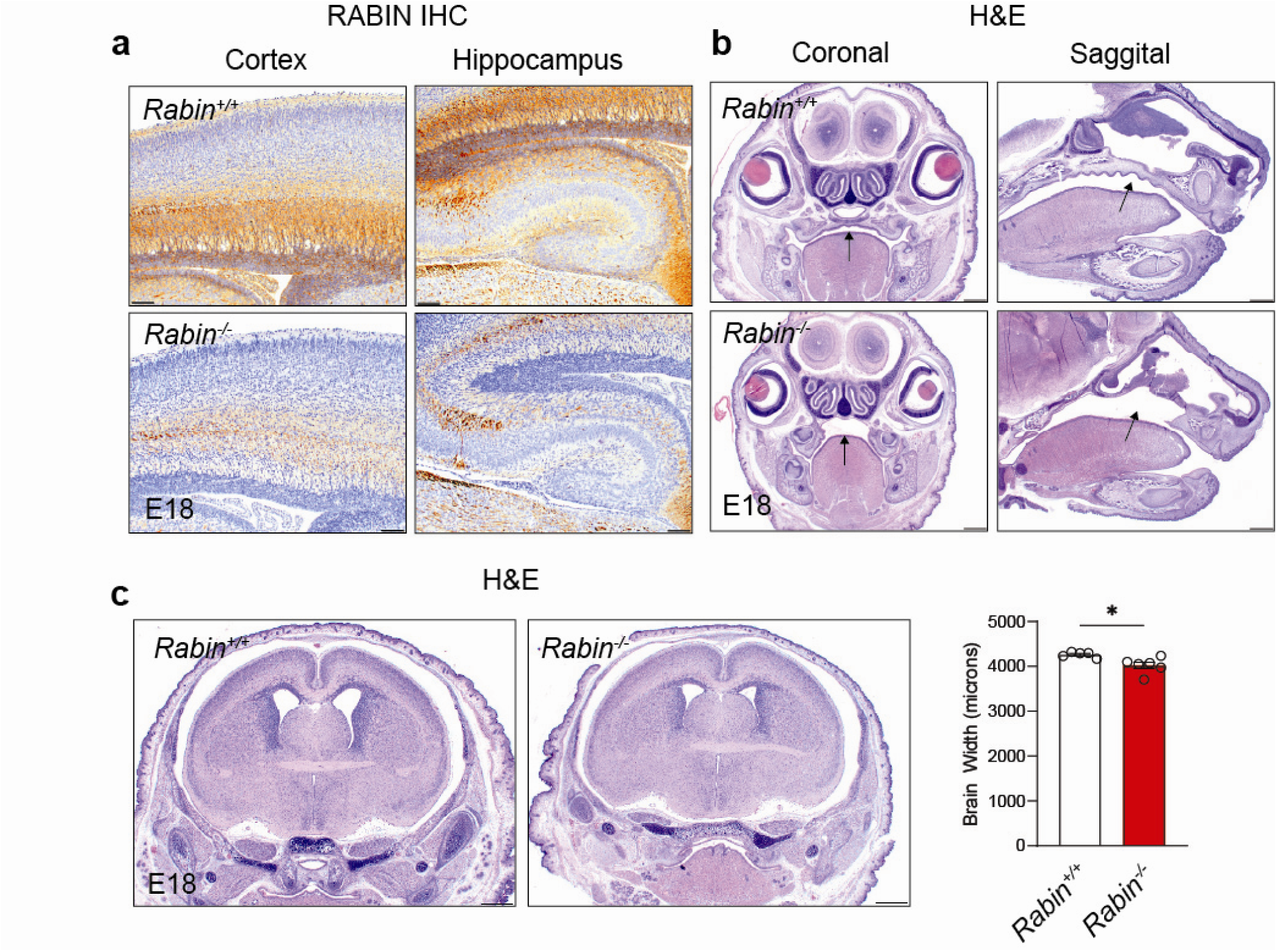
Craniofacial and whole-brain defects in germline *Rabin^−/−^* mice. **a.** Neocortical and hippocampal anatomy in E18 *Rabin^−/−^* mouse embryos. Scale bars, 100 μm. **b.** All *Rabin^−/−^*embryos examined had a palate defect (arrows). **c.** *Rabin^−/−^*embryos also had smaller brains than their littermate *Rabin^+/+^*controls, as measured by total brain width. Welch’s two-tailed *t*-test, *t*=2.86, \**p*=0.0188, n=5, 6 mice. Averaged data are presented as the mean ± SEM.

**Supplementary Figure S8.**
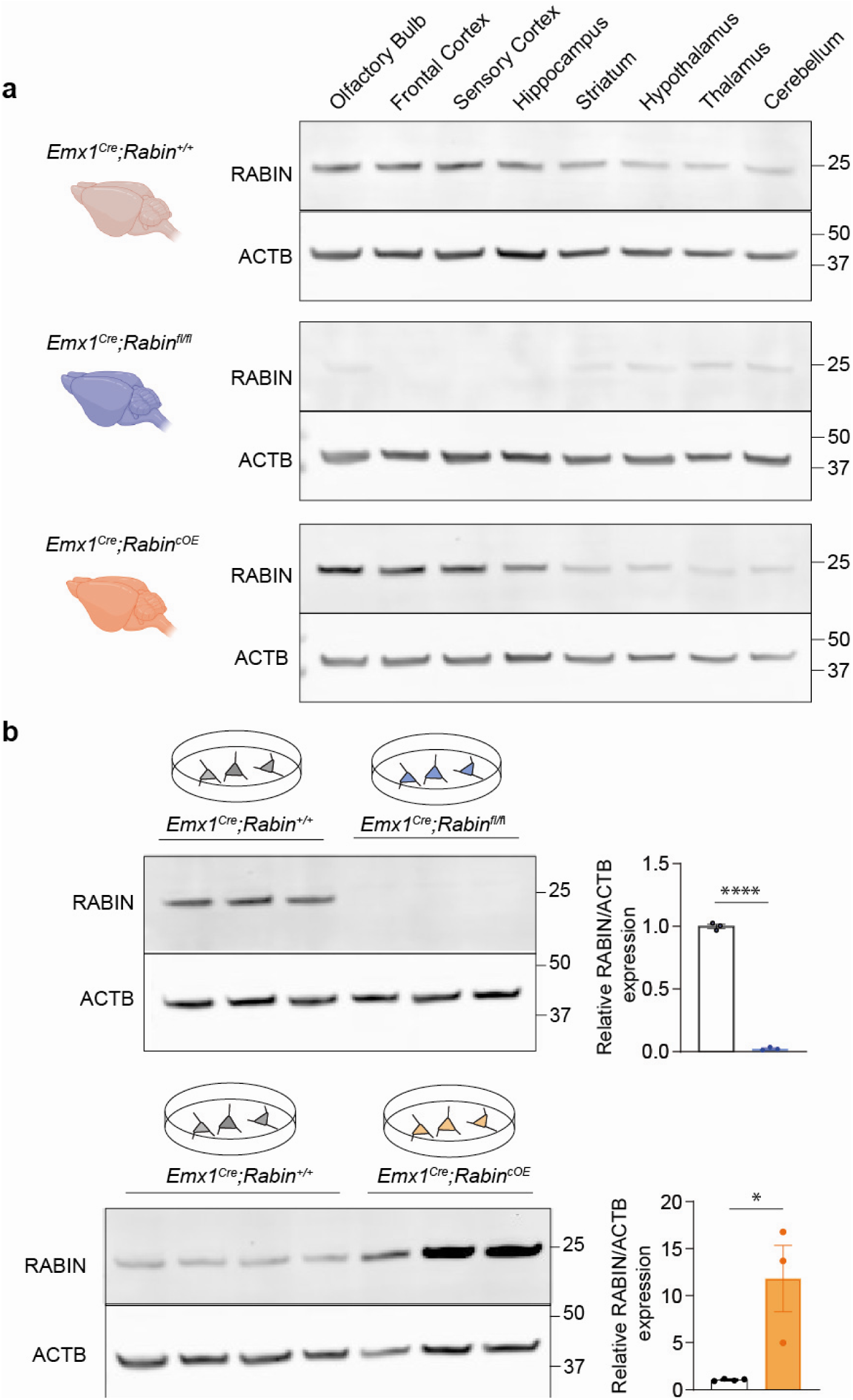
Validation of *Emx1^Cre^;Rabin^fl/fl^* conditional-knockout and *Emx1^Cre^;Rabin^OE^* overexpression mice. **a.** Confirmation of forebrain-specific conditional RABIN protein expression. **b.** Primary hippocampal neurons from *Emx1^Cre^;Rabin^fl/fl^* and *Emx1^Cre^;Rabin^cOE^*mice confirm conditional loss or overexpression of RABIN relative to littermate controls. *Emx1^Cre^;Rabin^fl/fl^*: Welch’s one-tailed *t*-test, *t*=54.30, \*\*\*\**p*<0.0001, n=3, 3 littermate cultures. *Emx1^Cre^;Rabin^cOE^:* Welch’s one-tailed *t*-test, *t*=3.050, \**p*=0.0464, n=4, 3 littermate cultures. Averaged data are presented as the mean ± SEM.

**Supplementary Figure S9.**
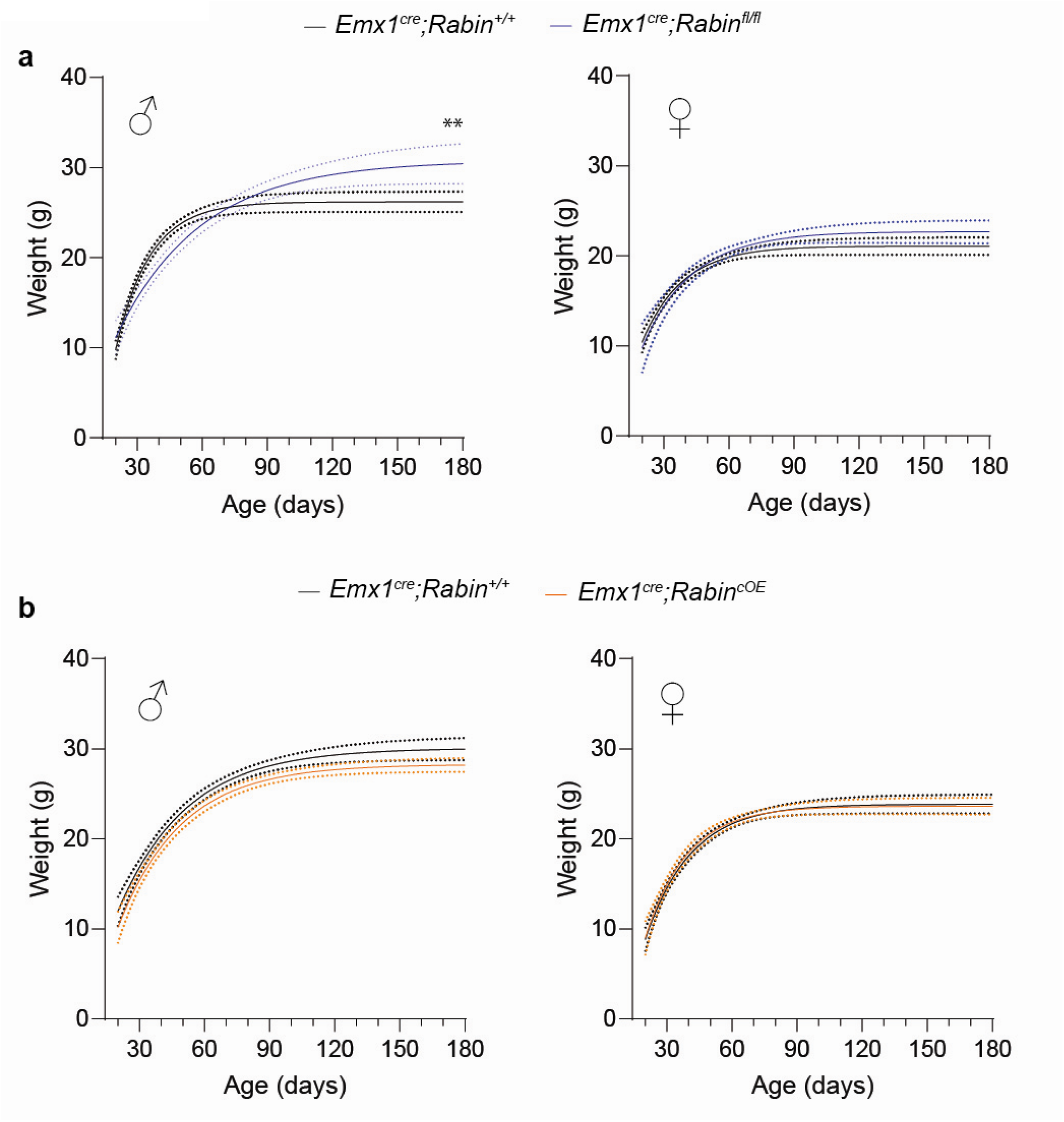
Single exponential fits to the growth curves of *Emx1^Cre^*;*Rabin^fl/fl^*mice and *Emx1^Cre^*;*Rabin^cOE^* mice. **a.** Male, but not female, *Emx1^Cre^;Rabin^fl/fl^* mice develop greater body mass relative to littermate controls. Males: Welch’s two-tailed *t*-test, *t*=2.991, \*\**p*=0.0041, n=51, 42 mice. Females: Welch’s two-tailed *t*-test, *t*=1.657, *p*=0.1008, n=67, 44 mice. **b.** The body mass of male and female *Emx1^Cre^;Rabin^cOE^* mice are indistinguishable from littermate controls. Males: Welch’s two-tailed *t*-test, *t*=1.801, *p*=0.0736, n=111, 127 mice. Females: Welch’s two-tailed *t*-test, *t*=0.2488, *p*=0.8038, n=63, 103 mice.

**Supplementary Figure S10.**
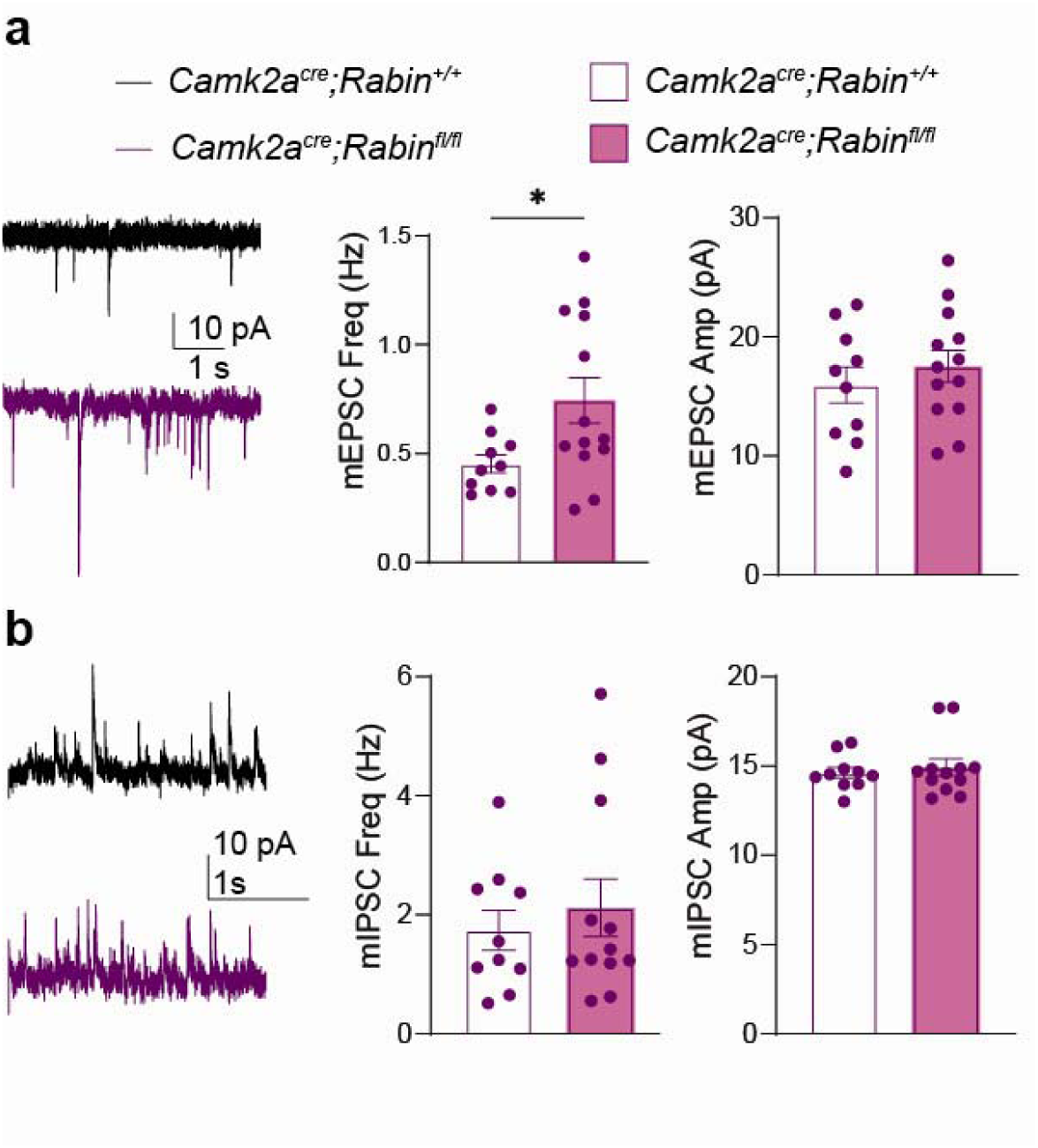
*Camk2a^Cre^*-mediated *Rabin* deletion increases the frequency of miniature excitatory postsynaptic currents. **a.** (Left) Representative mEPSC traces in *Camk2a^Cre^;Rabin^fl/fl^*mice (magenta) and littermate controls (black). (Middle) The mEPSC frequency is increased in CA1 pyramidal neurons with *Camk2a^Cre^*-mediated *Rabin* deletion. Welch’s two-tailed *t*-test, *t*=2.588, \**p*=0.0201, n=10, 13 cells. (Right) The mEPSC amplitude is unchanged between *Camk2a^Cre^;Rabin^fl/fl^* and littermate controls. Welch’s two-tailed *t*-test, *t*=0.7834, *p*=0.4428, n=10, 13 cells. **B.** (Left) Representative miniature inhibitory postsynaptic current (mIPSC) traces in *Camk2a^Cre^;Rabin^fl/fl^*(magenta) mice and littermate controls (black). (Middle) CA1 pyramidal neuron mIPSC frequency is indistinguishable between *Camk2a^Cre^;Rabin^fl/fl^*mice and controls. Welch’s two-tailed *t*-test, *t*=0.6366, *p*=0.5321, n=10, 12 cells. (Right) The mIPSC amplitude is unchanged between *Camk2a^Cre^;Rabin^fl/fl^*and littermate controls. Mann-Whitney *U* test, *U*=54.50, *p*=0.7351, n=10, 12 cells. Averaged data are presented as the mean ± SEM.

**Supplementary Figure S11.**
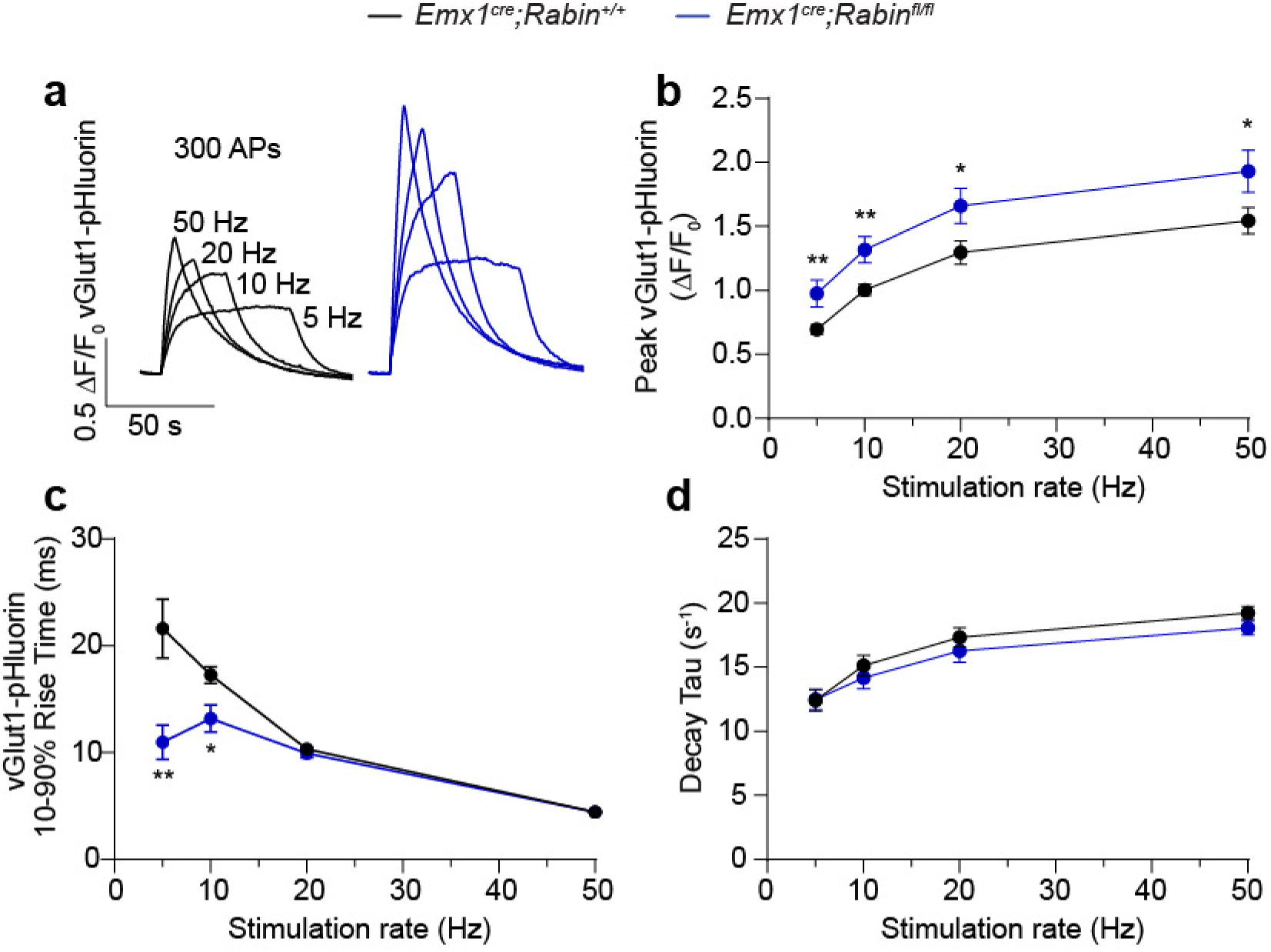
*Rabin* deletion enhances the magnitude and accelerates the kinetics of evoked vGlut1-pHluorin responses. **a.** Representative vGlut1-pHluorin fluorescence traces evoked by 300 stimuli delivered at 5, 10, 20, or 50 Hz in primary hippocampal cultures from littermate controls (black) and *Emx1^Cre^;Rabin^fl/fl^* (blue) mice. **b.** *Emx1^Cre^;Rabin^fl/fl^* synapses exhibited larger peak vGlut1-pHluorin responses than did littermate controls across stimulation frequencies. Multiple unpaired *t*-tests, 5 Hz: *t*=2.897, \*\**p*=0.0089 n=13, 9 FOVs; 10 Hz: *t* = 3.103, \*\**p*=0.0056 n=13, 9 FOVs; 20 Hz: *t*=2.260, \**p*=0.0351 n=13, 9 FOVs; 50 Hz: *t*=2.119, \**p*=0.0468 n=13, 9 FOVs. **c.** Onset kinetics of vGlut1-pHluorin responses were faster in *Emx1^Cre^;Rabin^fl/fl^*synapses at lower stimulation frequencies (5 and 10 Hz) but not at higher frequencies. Multiple unpaired *t*-tests, 5 Hz: *t*=3.050, \*\**p*=0.0063 n=13, 9 FOVs; 10 Hz: *t*=2.507, \**p*=0.0209 n=13, 9 FOVs; 20 Hz: *t*=1.018, *p*=0.3208 n=13, 9 FOVs; 50 Hz: *t*=0.2156, *p*=0.8314 n=13, 9 FOVs. **d.** Vesicle re-acidification kinetics, assessed by vGlu1-pHluorin decay, were indistinguishable between genotypes across all stimulation frequencies. Multiple unpaired *t*-tests, 5 Hz: *t*=0.2005, *p*=0.8431, n=13, 9 FOVs; 10 Hz: *t*=0.5085, *p*=0.6167, n=13, 9 FOVs; 20 Hz: *t*=0.5387, *p*=0.5961, n=13, 9 FOVs; 50 Hz: *t*=1.466, *p*=0.1581, n=13, 9 FOVs. Averaged data are presented as the mean ± SEM.

**Supplementary Figure S12.**
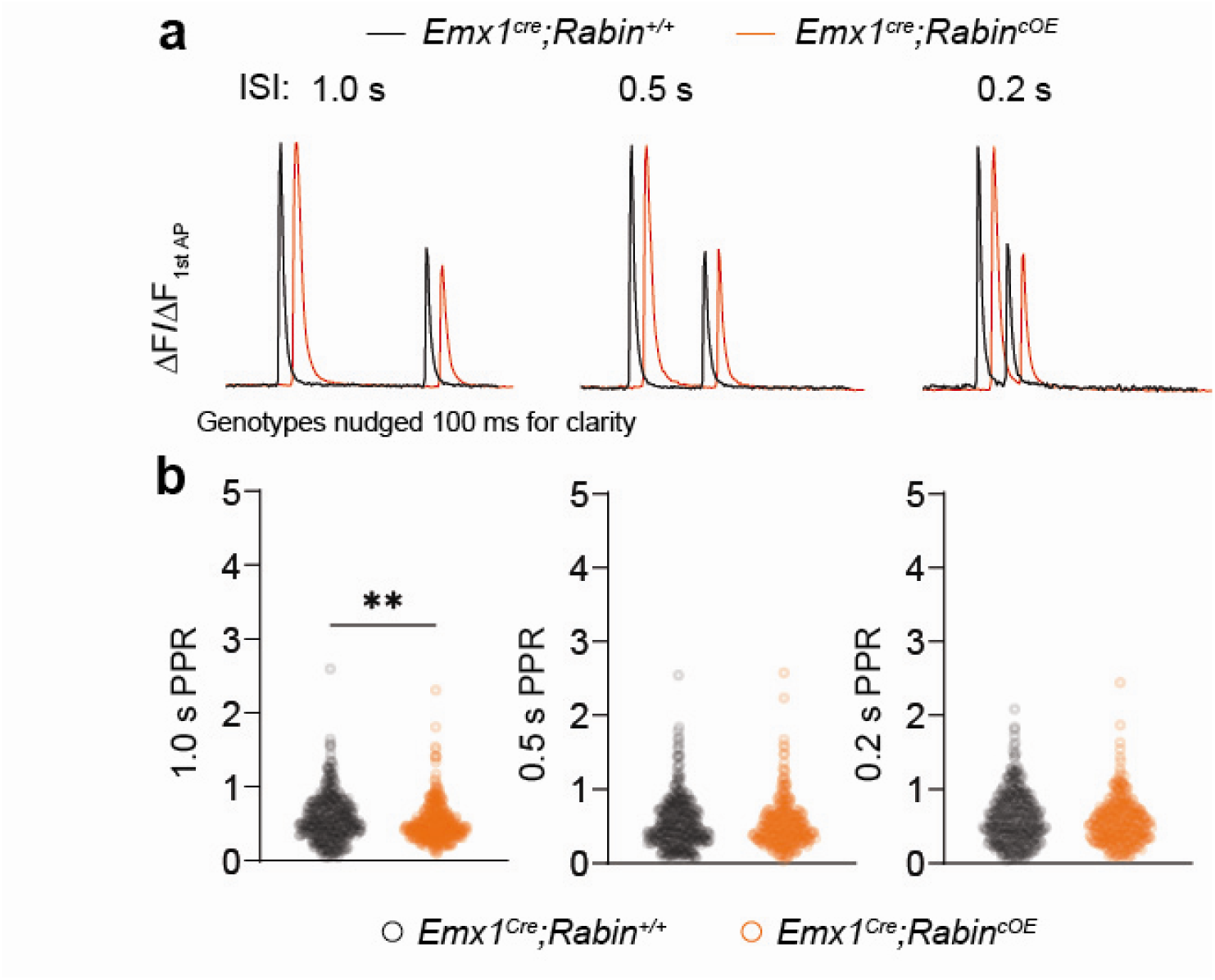
*Emx1^Cre^;Rabin^cOE^*synapses have greater paired-pulse depression (PPD) than those of littermate controls. **A.** Representative traces of iGluSnFr synaptic responses to pairs of field stimuli 1.0, 0.5, or 0.2 s apart in *Emx1^Cre^;Rabin^cOE^*(orange) or littermate control (black) neurons. B. *Emx1^Cre^;Rabin^cOE^*synapses have greater PPD than do littermate control neurons at the 1.0-s ISI but not at the 0.5-s or 0.2-s ISI. 1.0-s ISI: Mann-Whitney *U* test, *U*=63379, \*\**p*=0.0022, n=394, 369 synapses. 0.5-s ISI: Mann-Whitney *U* test, *U*=55882, *p*=0.5992, n=374, 306 synapses. 0.2-s ISI: Mann-Whitney *U* test, *U*=61275, *p*=0.1250, n=377, 348 synapses.

**Supplementary Figure S13.**
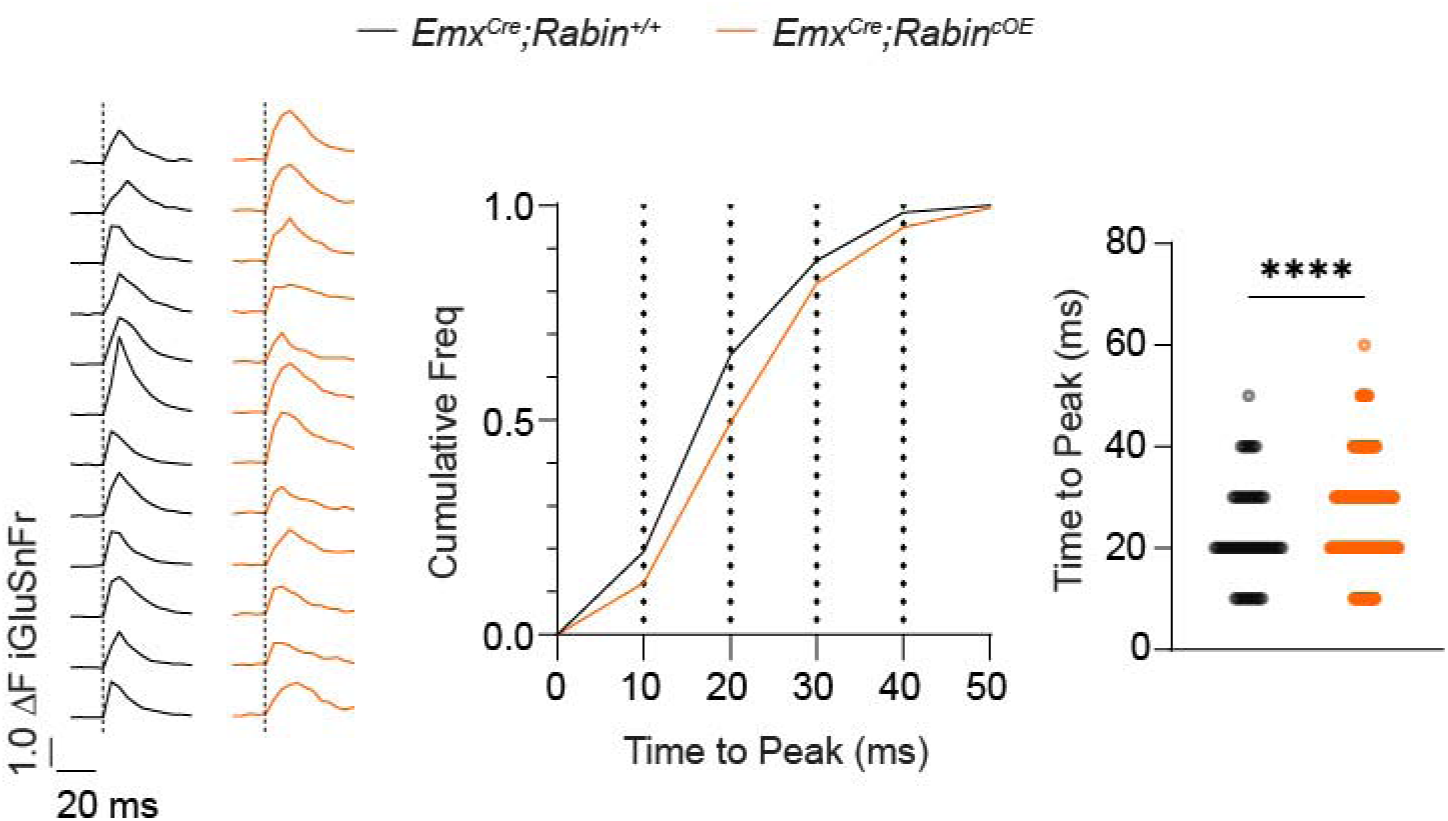
*Emx1^Cre^;Rabin^cOE^*synapses displayed slower evoked iGluSnFr kinetics than that in littermate controls. (Left) Representative iGluSnFr fluorescence traces measured at single synapses immediately after a field stimulus (indicated by the dashed vertical lines) for each indicated genotype. (Middle) Cumulative distribution plot of the time-to-peak fluorescence for all synapses. (Right) Time-to-peak fluorescence is plotted for each synapse and compared between the indicated genotypes. Mann-Whitney *U* test, *U*=151175, \*\*\*\**p* <0.0001, n=454, 809 synapses.

**Supplementary Figure S14.**
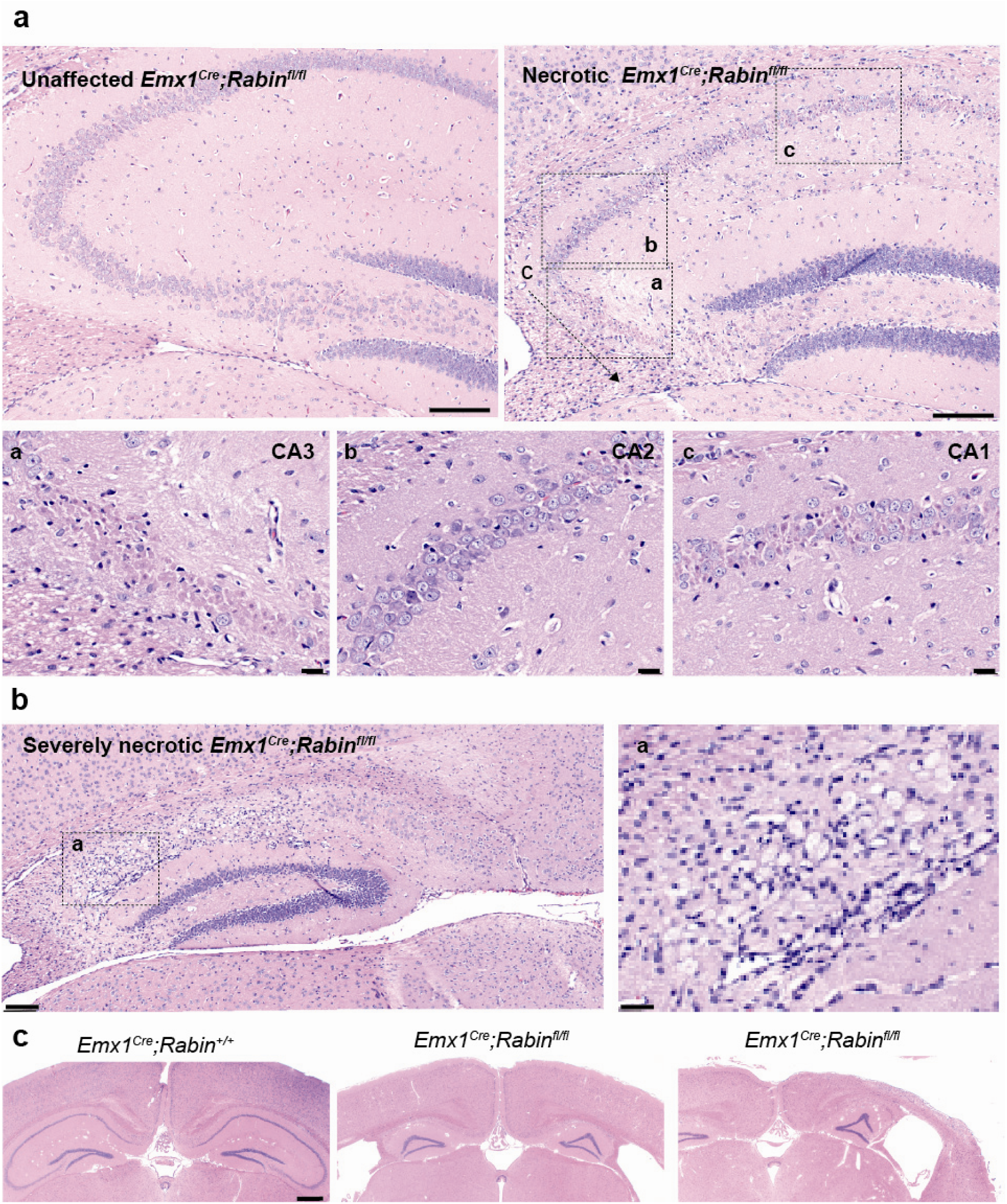
Hippocampal and cortical sclerosis is evident in a subset of *Emx1^Cre^;Rabin^fl/fl^*mice. **a.** Representative H&E-stained hippocampal sections from *Emx1^Cre^;Rabin^fl/fl^*mice that were either unaffected (left) or had necrotic tissues (right and bottom panels). Within the hippocampus, sclerosis is most common in areas CA3 (a) and CA1 (c) but not in CA2 (b). Scale bars: 100 μm. Insets: Scale bars, 20 μm. **b, c.** In the most severe cases, sclerosis extended across hippocampal CA1–CA3 areas and the neocortex. **b**. Scale bars, 100 μm. **c**., Scale bar, 500 μm.

**Supplementary Figure S15.**
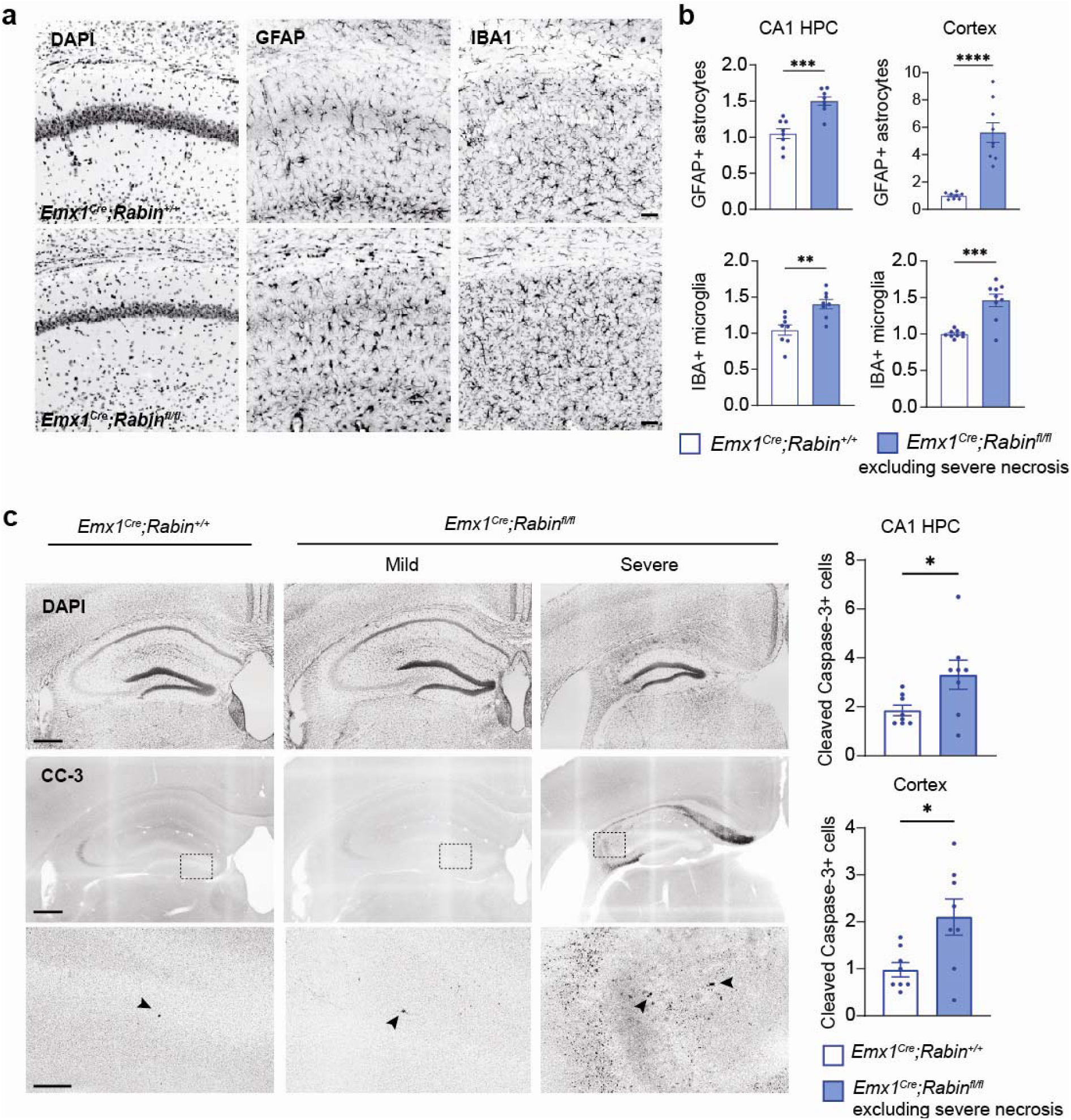
The *Emx1^Cre^;Rabin^fl/fl^* genotype results in increased astrocyte and microglia cell density with increased expression of cleaved caspase-3 in the hippocampus and neocortex. **a.** Representative micrographs of the CA1 region of the hippocampus (HPC) stained for DAPI and immunolabeled for cell markers of astrocytes (GFAP) and microglia (IBA1). Scale bars, 50 μm. **b.** Quantification of the density of astrocytes and microglia in the CA1 region revealed increased inflammatory cell markers in *Emx1^Cre^;Rabin^fl/fl^* mice. CA1 astrocytes: Welch’s two-tailed *t*-test, *t*=4.957, \*\*\**p* <0.0002, n=8, 8 mice. CA1 microglia: Welch’s two-tailed *t*-test, *t*=3.701, \*\**p* <0.0024, n=8, 8 mice. Cortex astrocytes: Welch’s two-tailed *t*-test, *t*=6.021, \*\*\*\**p* <0.0001, n=8, 9 mice. Cortex microglia: Welch’s two-tailed *t*-test, *t*=5.199, \*\*\**p* <0.0006, n=8,9 mice. **c.** Cleaved-caspase 3 (CC-3) staining indicated elevated apoptosis in *Emx1^Cre^;Rabin^fl/fl^*mice. Scale bars, 400 μm. Insets, Scale bar, 100 μm. CA1 CC-3: Welch’s two-tailed *t*-test, *t*=2.313, \**p*=0.0467, n=8, 8. Cortex CC-3: Welch’s two-tailed *t*-test, *t*=2.713, \**p*=0.0236, n=8, 8 mice. Averaged data are presented as the mean ± SEM.

**Supplementary Figure S16.**
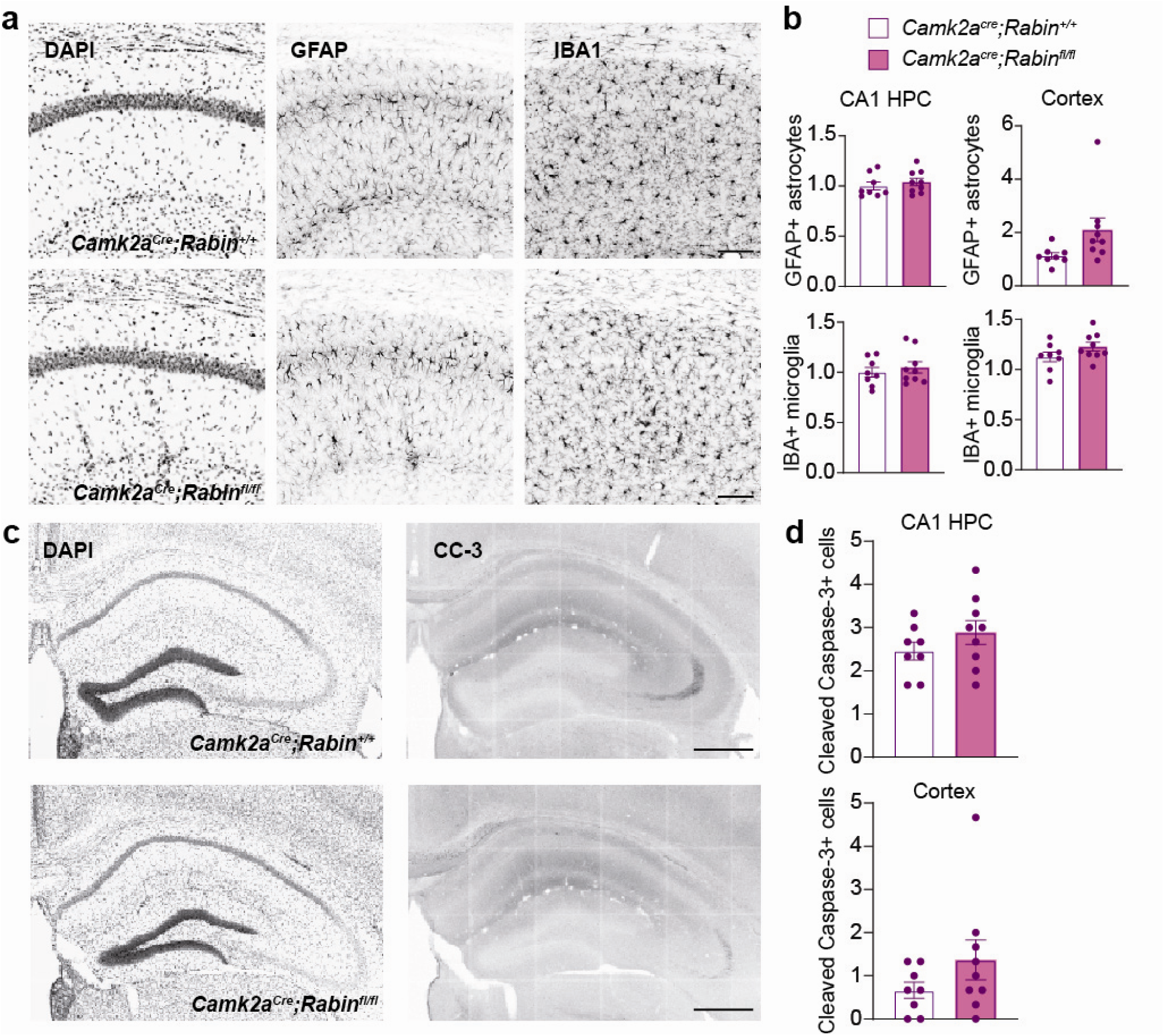
*Camk2a^Cre^;Rabin^fl/fl^*mice show no increase in astrocytes, microglia, or cleaved caspase-3 in the hippocampus or neocortex. **a.** Representative micrographs of the CA1 region of the hippocampus (HPC) stained for DAPI and immunolabeled for cell markers of astrocytes (GFAP) and microglia (IBA1). Scale bars, 100 μm. **b.** Quantification of the density of astrocytes and microglia in the CA1 region in *Camk2a^Cre^;Rabin^fl/fl^*mice. CA1 astrocytes: Welch’s two-tailed *t*-test, *t*=0.7145, *p*=0.4861, n=8, 9 mice. CA1 microglia: Welch’s two-tailed *t*-test, *t*=0.6439, *p*=0.5294, n=8, 9 mice. Cortex astrocytes: Welch’s two-tailed *t*-test, *t*=2.141, *p*=0.0605, n=8, 9 mice. Cortex microglia: Welch’s two-tailed *t*-test, *t*=1.555, *p*=0.1413, n=8, 9. **c.** Cleaved-caspase 3 (CC-3) staining was unchanged in *Camk2a^Cre^;Rabin^fl/fl^* mice. Scale bars, 500 μm. CA1 CC-3: Welch’s two-tailed *t*-test, *t*=1.240, *p*=0.2349, n=8, 9 mice. Cortex CC-3: Welch’s two-tailed *t*-test, *t*=1.407, *p*=0.1881, n=8, 9 mice. Averaged data are presented as the mean ± SEM.

**Supplementary Figure S17.**
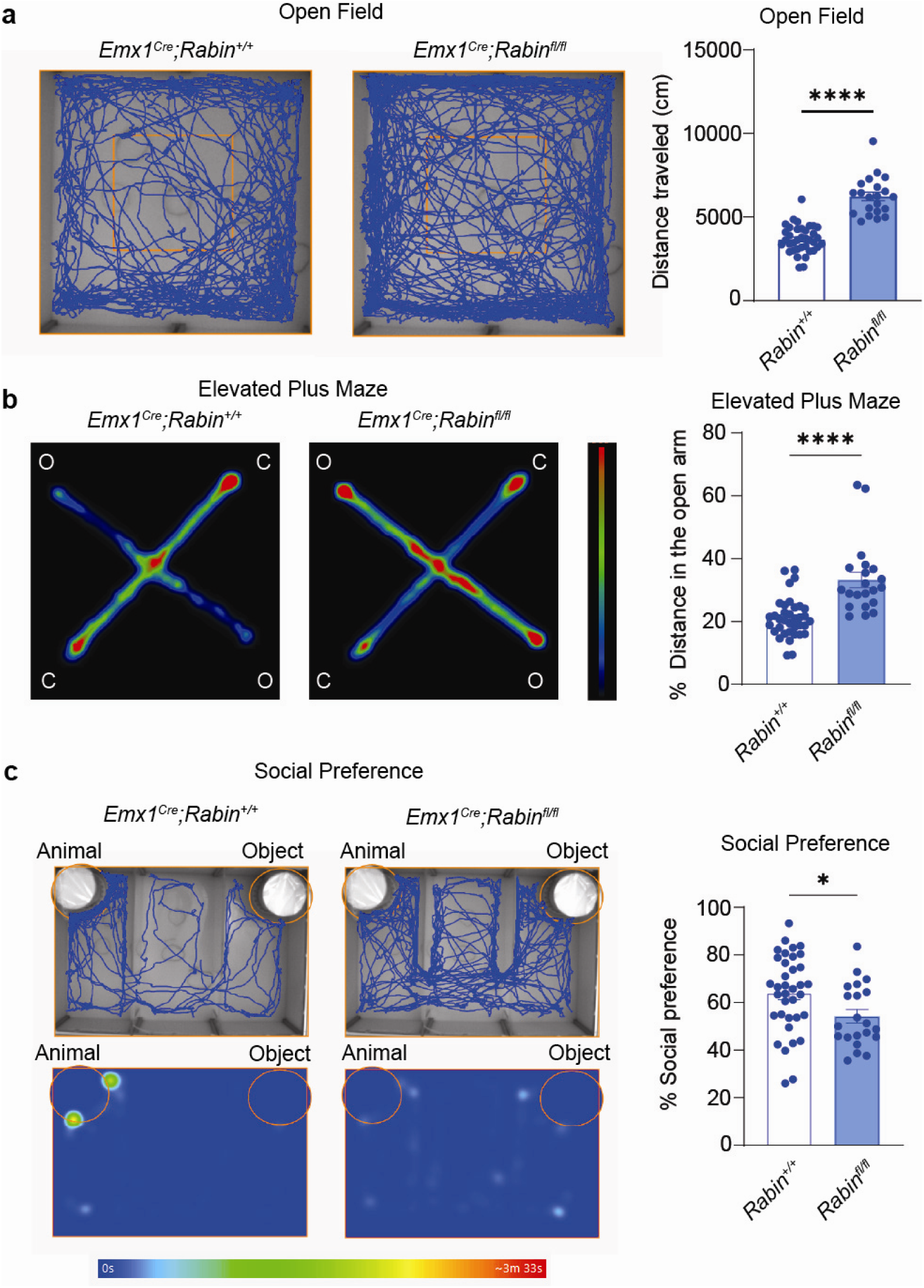
*Emx1^Cre^;Rabin^fl/fl^*mice exhibit epilepsy-associated behavioral deficits. **a.** *Emx1^Cre^;Rabin^fl/fl^*mice showed increased open-field locomotion, compared to littermate controls. Welch’s two-tailed *t*-test, *t*=8.988, \*\*\*\**p* <0.0001, n=37, 21 mice. **b.** *Emx1^Cre^;Rabin^fl/fl^*mice also exhibited increased open-arm exploration in the elevated plus maze. Welch’s two-tailed *t*-test, *t*=4.550, \*\*\*\**p* <0.0001, n=36, 21 mice. **c.** *Emx1^Cre^;Rabin^fl/fl^* mice showed deficits in normal social behavior. Welch’s two-tailed *t*-test, *t*=2.458, \**p*=0.0175, n=36, 21 mice. Averaged data are presented as the mean ± SEM.

**Supplementary Figure S18.**
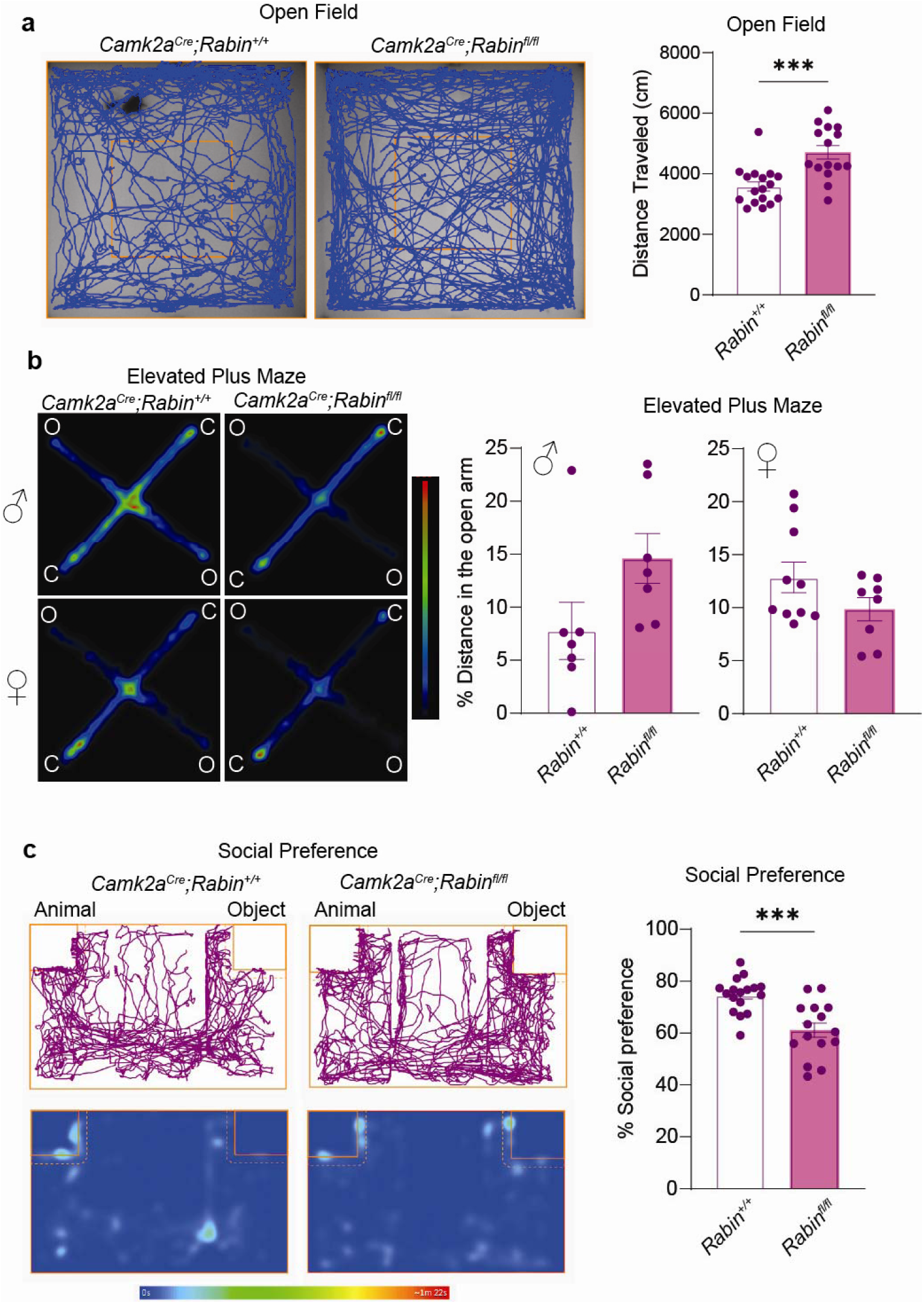
*Camk2a^Cre^;Rabin^fl/fl^*mice exhibit epilepsy-associated behavioral deficits. **a.** *Camk2a^Cre^;Rabin^fl/fl^*mice showed increased open-field locomotion, compared to their littermate controls. Welch’s two-tailed *t*-test, *t*=4.156, \*\*\**p*=0.0003, n=17, 15 mice. **b.** *Camk2a^Cre^;Rabin^fl/fl^* mice exhibited similar open-arm exploration in the elevated plus maze, compared to littermate controls. Males: Welch’s two-tailed *t*-test, *t*=1.902, *p*=0.0818, n=7, 7 mice. Females: Welch’s two-tailed *t*-test, *t*=1.655, *p*=0.1179, n=10, 8 mice. **c.** *Camk2a^Cre^;Rabin^fl/fl^* mice showed deficits in normal social behavior. Welch’s two-tailed *t*-test, *t*=4.237, \*\*\**p*=0.0003, n=17, 15 mice. Averaged data are presented as the mean ± SEM.

**Supplementary Figure S19.**
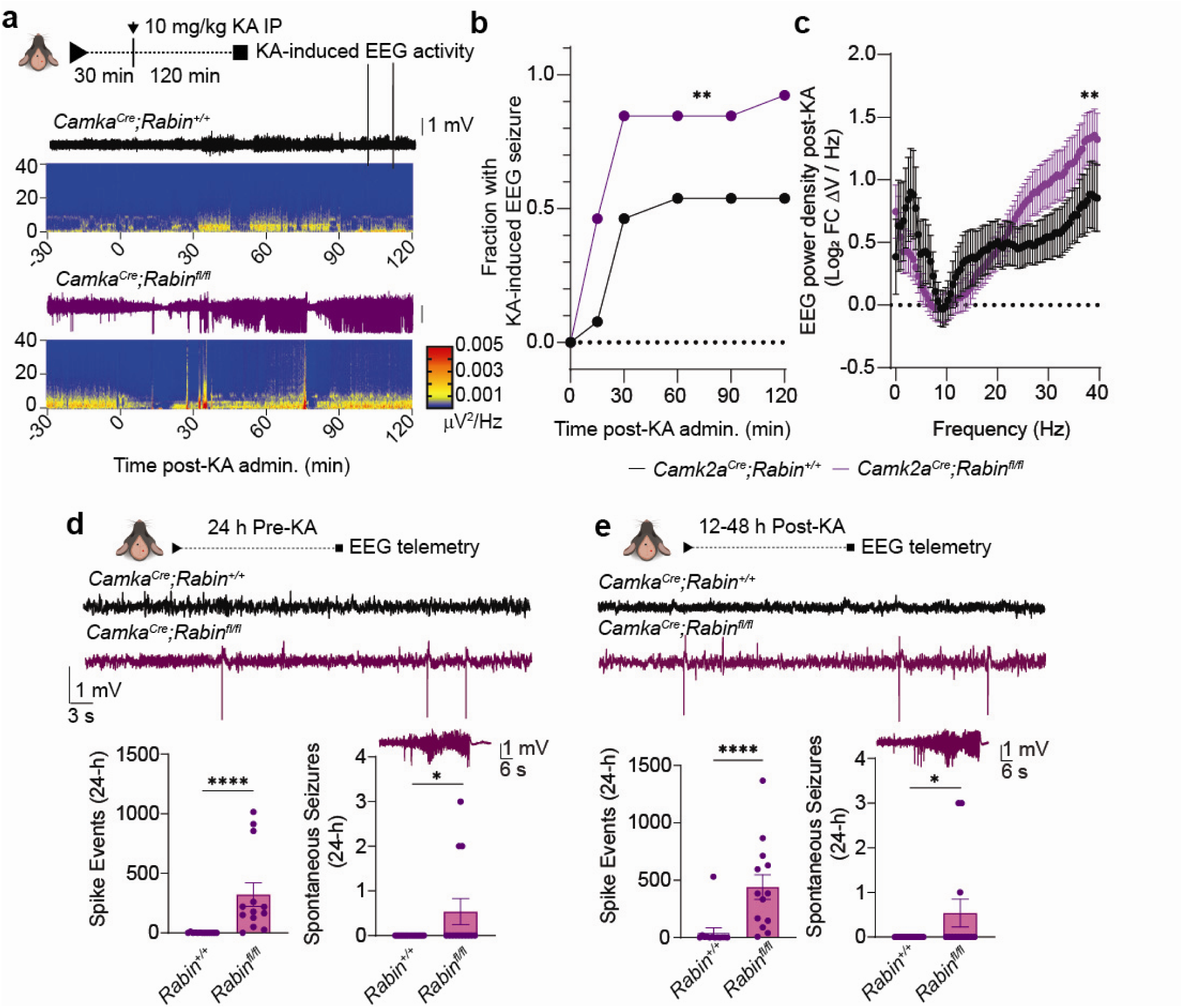
Epileptiform discharges and seizures in *Camk2a^Cre^;Rabin^fl/fl^* mice. **a.** EEG telemetry revealed an enhanced increase in EEG power in *Camk2a^Cre^*;*Rabin^fl/fl^* mice following a subthreshold intraperitoneal dose of kainic acid (KA). Representative EEG traces and corresponding power spectra are shown for control and *Camk2a^Cre^*;*Rabin^fl/fl^* mice for the 30 min preceding and 120 minutes following injection. **b**. *Camk2a^Cre^*;*Rabin^fl/fl^*mice exhibited significantly greater EEG seizure activity in response to KA, relative to littermate controls. Two-way ANOVA revealed a significant effect of genotype *F*(1,5)=23.41, \*\**p*=0.0054; n=13, 13 mice. **c**. EEG power was more strongly affected by a subthreshold dose of KA in *Camk2a^Cre^*;*Rabin^fl/fl^*mice. Mean Log_2_ fold change ± SEM. Two-way ANOVA revealed a significant effect of genotype: *F*(1,1886)=8.088, \*\**p*=0.0045; n=13, 13 mice. **d**. *Camk2a^Cre^*;*Rabin^fl/fl^*mice exhibited spontaneous epileptiform discharges in the 24-hour period prior to KA administration. Representative EEG traces are shown for control (black) and *Camk2a^Cre^*;*Rabin^fl/fl^* mice (purple). Spike and seizure events are plotted as mean ± SEM. Spike events: Mann-Whitney *U* test, *U=*4, \*\*\*\**p*<0.0001, *n* = 13, 13 mice. Seizure events: Unpaired one-tailed *t*-test, \**p*=0.0384, *n* = 13, 13 mice. **e**. Following KA washout, *Camk2a^Cre^*;*Rabin^fl/fl^*mice had increased spontaneous epileptiform discharges and seizures compared to littermate controls. Representative EEG traces are shown as in **d**. Spike events: Mann-Whitney *U* test, *U=*10, \*\*\*\**p*<0.0001, n=13, 13 mice. Seizure events: Unpaired one-tailed *t*-test, \**p*=0.0488, *n* = 13, 13 mice. Averaged data are presented as the mean ± SEM.

Movie S1. Spontaneous epileptiform discharges and seizure in an *Emx1^Cre^;Rabin^fl/fl^* mouse.

Table S1. Prioritization search for highly conserved, brain-enriched genes of unknown function.

**Table S2.** Structural effects of knocking out or overexpressing *Rabin*.

| Metric | Genotype<sup>a</sup> |  |  |  |
| --- | --- | --- | --- | --- |
|  | *Emx1<sup>Cre</sup>;Rabin<sup>+/+</sup>* | *Emx1<sup>Cre</sup>;Rabin<sup>fl/fl</sup>* | *Emx1<sup>Cre</sup>;Rabin<sup>+/+</sup>* | *Emx1<sup>Cre</sup>;Rabin<sup>cOE</sup>* |
| Synapse length (μm) | 166.9 ± 3.189 | 172.5 ± 2.944 | 148.1 ± 2.946 | 158.5 ± 3.084 |
| Active zone vesicles | 2.3 ± 0.13 | 2.5 ± 0.12 | 2.3 ± 0.18 | 2.5 ± 0.16 |
| Vesicle density | 7.7E<sup>-4</sup> ± 2.6E<sup>-5</sup> | 7.8E<sup>-4</sup> ± 1.9E<sup>-5</sup> | 1.0E<sup>-3</sup> ± 4.7E<sup>-5</sup> | 8.9E<sup>-4</sup> ± 3.2E<sup>-5</sup>** |

<sup>a</sup> All data are presented as the mean ± SEM.
\*\**p*<0.01

See the attached Excel sheet for all Supplemental Tables.

## MATERIALS AND METHODS

### Animal Care and Use

Mice of both sexes were used for all experiments. C57BL6/J mice and *Emx1^Cre^* mice were purchased from the Jackson Laboratory (JAX). *Camk2a^Cre^* mice were a gift from Ioannis Dragatsis (University of Tennessee Health Science Center, Memphis, TN). The care and use of animals were reviewed and approved by the Institutional Animal Care and Use Committee at St. Jude Children’s Research Hospital (St. Jude).

### Mutant Mice

#### Generation of Lrrc57/Rabin germline and conditional knockout mice

The *Lrrc57/Rabin* germline knockout (KO) and conditional knockout (cKO) mouse models were created in the Center for Advanced Genome Engineering and the Genetically Engineered Mouse Model Shared Resource (St. Jude) by using CRISPR technology and direct zygote injection as previously described^52^. Briefly, a mixture of the sgRNAs, Cas9 protein, and ssODN (single-strand oligodeoxynucleotide) donor templates, consisting of 60 ng/μL 3× NLS (triple nuclear localization signal) *Sp*Cas9 protein (St. Jude Protein Production Core), 20 ng/μL of each sgRNA (Synthego), and 5-10 ng/μL of each ssODN donor (IDT) were injected into the pronucleus of fertilized oocytes. Resulting pups were genotyped via targeted amplicon sequencing and analyzed using CRIS.py (https://www.ncbi.nlm.nih.gov/pmc/articles/PMC6414496/). Animals positive for the desired mutation sites were backcrossed to C57BL/6J mice and then bred to homozygosity. Editing construct sequences and relevant primers are listed in **Table S3.**

**Table S3.**
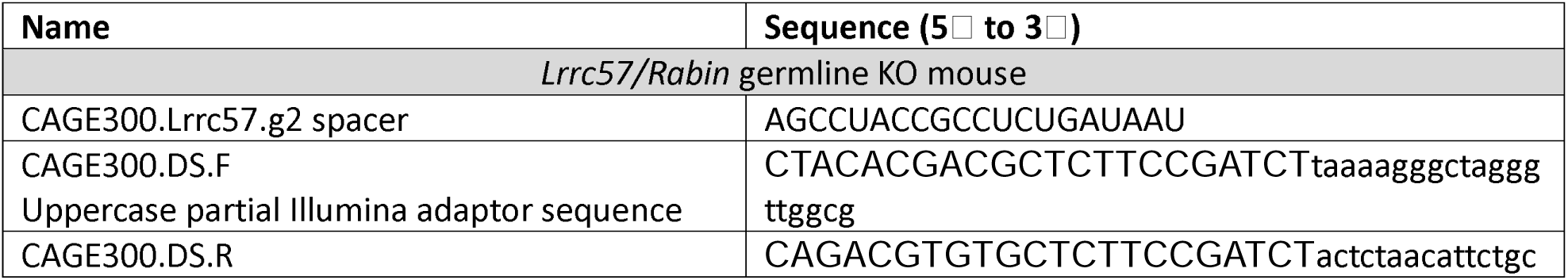
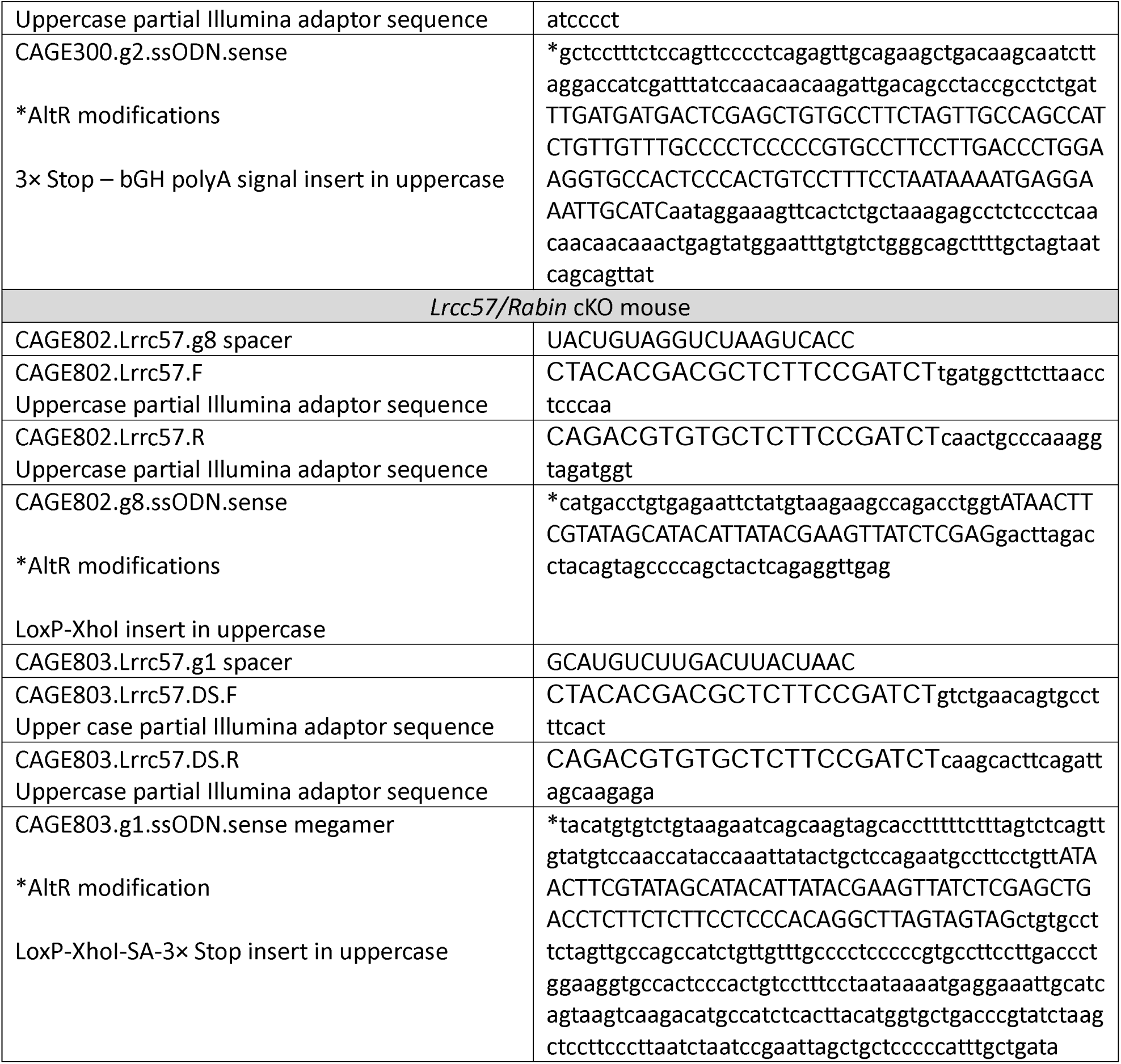
Construct sequences and primers used to generate *Lrrc57/Rabin* knockout mice.

#### Gene ration of Lrrc5 7/Ra bin-conditional overexpression allele

The cDNA of murine *Lrrc57/Rabin* was subcloned into a Transgenyx vector, such that the resulting transgene constructs consisted of SalI site–CAG promoter-chimeric intron-LoxP-small t intron-SV poly A signal-LoxP-Kozak sequence-Lrrc57/Rabin cDNA-IRES-mCherry-β-globin poly(A) signal-SalI site. The plasmid was linearized with SalI restriction enzyme (NEB R3138), and the resulting linear DNA was used for pronuclear injection into the male pronucleus of fertilized C57BL/6J oocytes. Founder mice were identified by PCR genotyping and maintained on the C57BL/6J genetic background.

### Generation of a Rabbit Monoclonal Antibody Against LRRC57/RABIN Protein

As a target peptide for the development of a monoclonal antibody, we chose a sequence of 12 amino acids (YMERFTATKKKF; **Supp. Fig. S2A**) from the C terminus of the protein because it is the most evolutionarily conserved part of the protein and because this sequence forms an exposed α-helix suitable as an antibody-binding site.

Four New Zealand White rabbits were immunized with the target peptide by using a standard protocol of five injections and two bleedings for each rabbit. Three subcutaneous injections were performed using the KLH immunogens, followed by two subcutaneous injections using the ovalbumin immunogens. For each immunization, the peptide conjugate was emulsified with complete Freund’s adjuvant for the primary injection and with incomplete Freund’s adjuvant for subsequent boosts. Serum samples (25 mL) were collected after the fourth and fifth immunizations (50 mL total), and antibodies were affinity-purified and screened in parallel.

The rabbit producing the most specific response was selected for monoclonal antibody production and was intravenously boosted with immunogen 4 days before splenectomy. A hybridoma was generated as described previously^53^. Briefly, splenocytes were fused with rabbit plasmacytoma cells (240E-W2) by using PEG4000 (Sigma PHR2892), and hybridomas were selected in HAT medium. Expanding clones from the initial 96-well plates were transferred to 24-well plates for continued culture. Supernatants were screened by Western blotting, and one clone (designated 1A1) displaying minimal background and high antigen specificity was selected.

Heavy- and light-chain variable regions containing complementarity-determining regions (CDRs) were sequenced and subcloned into the pLVX-EF1α-puro expression vector. Lentiviral particles were generated and used to transduce FreeStyle 293T cells (Invitrogen K900001). The resulting stable line (293T-1A1) was maintained in FreeStyle 293 serum-free medium (Thermo Fisher 12338018). Culture supernatants were collected after 7 days, filtered, and purified using a 1-mL Protein G affinity column (Cytiva Life Sciences). Bound antibody was eluted with 0.2 M glycine (pH 2.5) in a linear gradient and immediately neutralized with 1 M Tris (pH 9.0).

### Bioinformatic Analysis

To identify deeply conserved proteins of unknown function across Metazoan species, we began with a curated set of 1862 human proteins of unknown function, as compiled by Duek et al^54^. Proteins from this list were queried using the BLASTP algorithm against all available Metazoan non-Chordate proteomes in an all-against-all approach, using the Metazoa dataset (taxid: 33208), while excluding Chordata (taxid: 7711). A threshold of 75% biochemical similarity was applied using the BLOSUM62 scoring matrix without using sequence identities. Finally, the biochemical similarity between each pair of homologs was calculated from one-to-one pairwise alignments using Needleman-Wunsch algorithm (EMBOSS).The resulting subset of evolutionarily conserved proteins was then analyzed for enriched expression in the mouse brain by using data from ProteomicsDB and ranked using median protein expression values. From this brain-enriched subset of proteins of unknown function, we performed a PubMed search to identify any proteins whose functional annotation had been determined since 2019. These proteins were retained in the final list, and their reported functional annotations are presented in **Table S1**.

### Western Blots

Organs were dissected from 2-month-old C57BL/6J mice. Tissues were lysed in RIPA buffer, and Western blotting was performed as described previously^55,56^. The following antibodies were used in a given concentration: Antibody 1A1 against LRRC57/RABIN was generated in this study (250 ng/ml), GluA1 – 182003 (Synaptic Systems), 1:1000; GluN1 – 114011 (Synaptic Systems), 1:1000; ActB – MA5-15739-D680 (Invitrogen), 1:10.000, Syt1 – m Ab 48 (DSHB), 1μg/ml; Homer1 – 160011 (Synaptic Systems), 1:1000; Rab3 - 15029-1-AP (Proteintech), 1:1000; Rab8a - 6975T (Cell Signaling), 1:1000; SV2A - AB_2315387 (DSHB), 1:1000; VGlut1 – 135302 (Synaptic Systems), 1:1000; anti-HA (unpublished) at 100 ng/ml; anti-FLAG^57^ at 100 ng/ml; Rabbit anti-Goat – A27014 (Invitrogen) at 1:10000; Goat anti-rabbit – A27036 (Invitrogen) at 1:10000 and Goat anti-mouse – A28177 (Invitrogen) at 1:10000.

### Co-Immunoprecipitation

*Rabin/Lrrc57* cDNA was HA-tagged, whereas *Rab3a* and *Rab8a* cDNAs were FLAG-tagged and subcloned into lentiviral vectors. Embryonic day (E) 18.5 rat cortical neurons were maintained in culture as previously described^58^. Neurons were transduced with lentiviruses and lysed 48 h later in co-immunoprecipitation (co-IP) buffer (15 mM HEPES pH 7.4, 150 mM NaCl, 15 mM KCl, 1.5 mM MgCl₂, 0.15% IGEPAL CA-630).

Dynabeads™ M-270 Epoxy (Invitrogen; 14301) were coupled to IgG (Sigma I5506), anti-HA (unpublished data), or anti-FLAG^59^ antibodies prepared in house, per the manufacturer’s protocol. Cell lysates were supplemented with 20 mM EDTA, mixed, and incubated with either 200 µM GTPγS (Sigma G8634) or GDP (Sigma; G7127) at 30 °C for 30 min. MgCl₂ was added to 60 mM final concentration, and samples were incubated with antibody-coupled beads for 2 h at 4°C with gentle rotation. Beads were washed three times in high-stringency buffer (10 mM HEPES pH 7.4, 10 mM KCl, 50 mM NaCl, 1 mM MgCl₂, 0.05% IGEPAL CA-630) and once in low-stringency buffer (10 mM HEPES pH 7.4, 10 mM KCl, 0.07% IGEPAL CA-630). Bound proteins were eluted in 54 µL elution buffer (200 mM glycine pH 2.5, 1% SDS) by shaking for 10 min at room temperature in a ThermoMixer (Eppendorf) and immediately neutralized with 18 µL 1 M Tris pH 8.0 (final pH of the samples was 7.4). Eluates were analyzed by Western blot^55^.

### Subcellular Fractionation

Synaptosomes were isolated from mouse hippocampi by using Syn-PER reagent (ThermoFisher 87793) per the manufacturer’s protocol. Whole-cell, cytosolic, and S2 fractions were analyzed by Western blot.

### Isolation of Synaptic Vesicles

#### Tissue homogenization

Whole brains from 3-month-old male C57BL/6J mice were rapidly dissected and placed in 3 mL ice-cold Hibernate™-A medium (A1247501, Gibco). Each brain was then transferred into a Teflon-glass homogenizer containing 4 mL homogenization buffer (25 mM KCl, 25 mM consisting of KH₂PO₄ and K₂HPO₄, 5 mM EGTA, and protease inhibitors; cOmplete Mini EDTA-free, 1 tablet per 10 mL; pH 7.4) and kept on ice. Tissue homogenization was performed using an LS overhead stirrer (VELP Scientific, Inc.) at 900 rpm for 10 strokes. The resulting lysate was distributed into 2-mL Eppendorf tubes and centrifuged at 25,000 ×*g* for 20 min at 2 °C. The supernatant was carefully collected, and the KCl concentration was subsequently adjusted to 125 mM to achieve isotonic conditions^60^.

#### Antibody–bead conjugation

Antibody coupling to magnetic beads was performed as previously described^60^ with minor modifications. Briefly, 15 mg epoxidated M-PVA E02 magnetic beads (CMG-216, Revvity) were washed with 1 mL KPBS (145 mM KCl, 10 mM potassium phosphate consisting of KH₂PO₄ and K₂HPO₄, pH 7.4) and resuspended in 300 µL of borate buffer (100 mM boric acid, 76 mM NaOH, pH 8.5). Either bovine serum IgG (450 µg; I5506, Sigma-Aldrich) or anti-SV2α antibody (450 µg; DSHB AB_2315387) was adjusted to 300 µL with borate buffer and combined with the beads, followed by the addition of 300 µL of 3 M ammonium sulfate in borate buffer. The mixture was incubated overnight at 37 °C with rotation. The next day, antibody–bead conjugates were collected on a magnetic stand and washed alternately three times with wash buffer I (500 mM NaCl, 50 mM ammonium acetate, pH 4.5) and wash buffer II (500 mM NaCl, 50 mM Tris-HCl, pH 8.0). Unreacted epoxy groups were blocked with 1 M ethanolamine (Sigma 411000) dissolved in EveryBlot blocking buffer (Bio-Rad 12010020) for 15 min, followed by incubation in 1% BSA (001000162, Jackson ImmunoResearch Laboratory, Inc.) dissolved in PBS (pH 7.4) for 30 min at room temperature. The antibody–bead conjugates were washed twice with KPBS and stored at 4 °C until use.

#### Synaptic vesicle immunoprecipitation

Synaptic vesicle (SV) immunoprecipitation was performed as previously described^60^. A 200-µL aliquot of brain lysate was reserved for total protein analysis, and the remaining lysate was divided equally into two tubes (1.9 mL each) for SV pulldown. Antibody-coupled magnetic beads (15 mg total) were resuspended in 200 µL KPBS, and 100 µL of the bead suspension was added to each lysate. The tubes were placed on ice within 50-mL conical tubes and rotated for 20 min in a cold room. SV-bound beads were then washed three times with 1 mL ice-cold KPBS by gentle pipetting. Beads from each half-brain sample were combined and eluted with 150 µL elution buffer (2% SDS, 25 mM Tris-HCl, pH 8.0) at 50 °C for 5 min. The eluates were stored at −80 °C for subsequent Western blot analysis.

### Purification of Recombinant Proteins

#### LRRC57/RABIN

The pFastBac1-6xHis-HA-GlySer-mmLrrc57 construct was transformed into DH10Bac *Escherichia coli* and used to generate bacmid DNA per the manufacturer’s protocol (ThermoFisher 10359016). The bacmid was used to transfect Sf9 insect cells in Gibco SF900 III serum-free media. The transfected cell supernatants were harvested after 4 days, amplified to P3 generation virus, and used to infect Sf9 cells in suspension culture. The culture was harvested at 72 h postinfection and cell pellet lysed by homogenization in MCAC-0 buffer (50 mM Tris, pH 7.5, 500 mM NaCl, 10% glycerol). The lysate was loaded on 5-mL Nickel Affinity column (Cytiva Life Sciences) and eluted using 1M imidazole in MCAC-0 linear gradient. The eluted fraction was dialyzed into the storage buffer (15 mM HEPES, 150 mM NaCl, 15 mM KCl, 1.5 mM MgCl_2_, 0.15% IPEGAL CA-360) and incubated with His-tagged TEV protease overnight to remove the 6×His tag. The untagged protein was then purified on a reverse nickel affinity column. The cleaved LRRC57 was concentrated and loaded on an S100 16/60 SEC column (Cytiva) equilibrated in the final buffer to separate monomer from dimer; each corresponding peak was pooled separately for further analysis.

#### MBP-Rim1α(Rab3BD)

The pDG-MBP(N)-RIM1α(Rab3-BD)-Strep II construct was transformed into *E. coli* strain KRX (Promega L3002) and plated onto Luria broth (LB) agar supplemented with ampicillin. For expression, cells were grown in LB media at 37 °C. At OD_600 nm_ ∼0.6, the temperature was reduced to 20 °C, and cells were induced with 0.1% rhamnose for 20 h. The cells were collected by centrifugation, then suspended in lysis buffer (100 mM Tris-Cl, pH 7.4, 500 mM NaCl, 1 mM DTT, 10% glycerol, 0.2 % Triton-X 100) and lysed by three passages through a Panda 2000 homogenizer (GEA). The cleared lysate was then applied to an amylose resin (NEB E8022S) column and washed with 30 column volumes of lysis buffer before eluting the protein in lysis buffer plus 10 mM maltose. Pooled fractions were then loaded onto a Strep-Tactin^®^ 4Flow^®^ high-capacity FPLC column (IBA 2-1258-001) equilibrated in 50 mM Tris-Cl, pH 8.5, 500 mM NaCl, 5 mM DTT, 10% glycerol, 0.2 % Triton-X 100, washed with the same buffer, and eluted with 10 mM biotin. Finally, protein was dialyzed into 125 mM Tris-Cl, pH 7.4, 150 ml NaCl, 1mM DTT, 1mM EDTA, 10% glycerol. MBP control was purified similarly.

#### Rab3a(del)-FLAG

The pDG-Rab3a(del)WT-FLAG construct was transformed into *E. coli* strain KRX cells, as described for MBP-Rim1α(Rab3BD). Cell pellets were suspended in a lysis buffer containing 20 mM Tris, pH 7.4, 500 mM NaCl, 1 mM EDTA, 1mM DTT, and 10% glycerol and lysed by passaging them three times through a Panda 2000 homogenizer. Clarified supernatant was then applied to a Ni-NTA column (Gold Biotech H-350-5) and washed with 30 column volumes of lysis buffer plus 25 mM imidazole. Protein was eluted with 250 mM imidazole in the same buffer. Pooled fractions were incubated at 4 °C overnight with His-TEV protease, then filtered and applied to a Sephacryl S-100 HR 16/600 column (Cytiva) in 20 mM Tris-Cl, pH 7.4, 25 mM NaCl, 4 mM EDTA, 1 mM DTT. Pooled fractions were concentrated to 1 mg/mL.

#### Rab8a(del)-FLAG

The pET25-Rab8a(del)WT-FLAG construct was transformed into Rosetta2/pLysS cells (EMD Millipore 71401-3). Expression of the protein was induced with 1 mM IPTG at 20 °C overnight. Cell pellets were suspended in a lysis buffer containing 20 mM Tris-Cl, pH 7.4, 500 mM NaCl, 1 mM EDTA, 1mM DTT, and 10% glycerol and passaged three times through a Panda 2000 homogenizer. Clarified supernatant was then applied to a Ni-NTA column (Gold Biotech H-350-5). After a 30-column volume wash with 25 mM imidazole in lysis buffer, protein was eluted in the same buffer with 250 mM imidazole. Pooled fractions were digested with thrombin protease overnight at 4 °C. They were then concentrated and applied to a Sephacryl S-100 HR 16/600 column (Cytiva) in 20 mM Tris-Cl, pH 7.4, 25 mM NaCl, 4 mM EDTA, 1 mM DTT. Pooled fractions were concentrated to 1 mg/mL.

#### GST-OCRL

The pCool-GST-OCRL (mouse 538-900aa)-FLAG construct was transformed into *E. coli* strain Rosetta2/pLysS (EMD Millipore 71401-3). Expression of the protein was induced with 1 mM IPTG at 16 °C overnight. Cell pellets were suspended in lysis buffer of 125 mM Tris-Cl pH 7.4, 150 mM NaCl, 1 mM DTT and lysed by homogenization. After centrifugation, the supernatant was applied to a GST-affinity column (Cytiva 17513101) and washed extensively with lysis buffer. Protein was eluted with 10 mM reduced glutathione (Sigma G6529) in the same buffer. Protein was then concentrated and applied to a Superdex S-200 pg 26/600 column in 125 mM Tris pH 7.4, 150 mM NaCl, 1 mM DTT, 1 mM EDTA. Glycerol was added to the pooled fractions to a final concentration of 25% (v/v). GST control was purified similarly.

### Pulldown Studies with Recombinant Proteins

#### Probing a direct interaction between MBP-Rim1α(Rab3BD) and Rab3(del)-FLAG

Amylose magnetic beads (NEB E8035S) were loaded with 20 μg of the recombinant MBP-Rim1α(Rab3BD) (307.7 pmol) or full-length MBP, per the manufacturer’s instructions. RAB3A(del)WT [13.17 μg, corresponding to 1:2 molar ratio, with regard to MBP-Rim1α(Rab3BD)] stored in 20 mM Tris-Cl, pH 7.4, 25 mM NaCl, 4 mM EDTA, 1 mM DTT buffer, was loaded with either GDP or GTPγS, following the procedure described in the co-immunoprecipitation section above. Nucleotide-loaded Rab3a(del)WT was then mixed with recombinant LRRC57/RABIN (33.2 μg, corresponding to 1:2 molar ratio with regard to RAB3A), stored in 15 mM HEPES pH 7.4, 150 mM NaCl, 15 mM KCl, 1.5 mM MgCl_2_, 0.15% IGEPAL CA-630. These mixtures were incubated at 4 °C for 2 h. Amylose magnetic beads pre-loaded with either full-length MBP or MBP-Rim1α(Rab3BD) were resuspended in the storage buffer (50 mM HEPES-KOH, pH 7.2, 150 mM NaCl, 1 mM MgCl_2_, 0.1% Triton X-100). Once the interaction between nucleotide-loaded Rab3a(del)WT and LRRC57/RABIN was completed, magnetic beads with pre-loaded MBP or MBP-Rim1α(Rab3BD) were placed in a high-strength magnetic rack (Biorad #1614916), storage buffer was removed, and previously prepared mixtures of LRRC57/RABIN and nucleotide-bound RAB3A(del)WT were added to the beads. These mixtures were further incubated at 4 °C for 2 h. Finally, once this incubation step was completed, beads were washed three times with 1 mL washing buffer (10 mM HEPES-KOH, pH 7.2, 150 mM NaCl, 2 mM MgCl_2_, and 0.2% Triton X-100), and proteins were eluted with 30 µL elution buffer (20 mM maltose, 200 mM NaCl, 20 mM Tris-HCl pH 7.4, 1 mM EDTA, 1 mM DTT). Samples were stored at –20 °C until they were analyzed by SDS-PAGE. Briefly, samples were run in Tris/glycine gels, as previously described^55^; gels were then stained with GelCode™ Blue Stain Reagent (ThermoFisher 24592) and photographed on the light box (Hall Productions BL1218).

#### Probing a direct interaction between GST-OCRL(538-900) and Rab8(del)-FLAG

Pierce glutathione magnetic agarose beads (Thermo Fisher Scientific, 78601) were pre-loaded with 20 μg (280 pmol) recombinant GST–OCRL (mouse 538–900)–FLAG or full-length GST according to the manufacturer’s protocol. Appropriate amount of RAB8A(del)WT protein (11.3 μg) corresponding to the molar ratio 1:2, with regard to GST-OCRL(538-900), stored in 20 mM Tris-Cl, pH 7.4, 25 mM NaCl, 4 mM EDTA, 1 mM DTT, was pre-loaded with either GDP or GTPγS, following the procedure described in the co-immunoprecipitation section above. Nucleotide-loaded RAB8A(del)WT samples were then combined with 30.2 μg recombinant LRRC57/RABIN (1:2 molar ratio with regard to RAB8A(del)WT) and incubated for an additional 2 h at 4 °C. Separately, glutathione magnetic beads loaded with full-length GST or GST–OCRL(538–900)–FLAG were resuspended in storage buffer (125 mM Tris, pH 7.4, 150 mM NaCl, 0.1% Tween-20, 1 mM DTT, 1 mM EDTA). After the incubation of nucleotide-bound RAB8A(del)WT and LRRC57/RABIN was completed, the beads were placed on a high-strength magnetic rack (Bio-Rad #1614916), and the storage buffer was removed. The pre-assembled RAB8A(del)WT-LRRC57/RABIN complexes were then added to the magnetic beads and incubated for an additional 2 h at 4 °C. Once the last incubation was completed, beads were washed three times with 1 mL washing buffer (identical to the storage buffer), and proteins were eluted in 30 µL elution buffer (50 mM reduced glutathione, 125 mM Tris, pH 8.1, 150 mM NaCl, 0.1% Tween-20, 1 mM DTT, 1 mM EDTA). Eluted samples were stored at –20 °C until analysis. Samples were resolved by SDS–PAGE by using Tris/glycine gels as previously described^4^, stained with GelCode™ Blue Stain Reagent (Thermo Fisher 24592), and visualized on a light box (Hall Productions BL1218).

### Electron Microscopy

#### Immuno-electron microscopy

For post-embedding immuno-electron microscopy, four WT mice were used. Animals were transcardially perfused with 4% paraformaldehyde (PFA) in 0.1 M Sørensen’s phosphate buffer (PB). The hippocampal CA3 region was isolated by vibratome sectioning followed by manual dissection in ice-cold PB to obtain tissue blocks of approximately 0.8 mm × 0.8 mm × 0.15 mm.

Tissue blocks were sandwiched between 3-mm copper planchettes (Type A, 200-µm cavity; Type B, flat), both coated with 1-hexadecene. The cavity of the Type A planchette was filled with 20% BSA to eliminate air gaps. Tissue samples were sandwiched within the planchettes and cryoimmobilized using a high-pressure freezer (Leica EM ICE, Leica Microsystems) at liquid nitrogen temperature, between 2040 – 2050 bars, with a cooling rate of ∼20,000 K/s, and a freezing time of ∼20 ms.

Frozen specimens were transferred to a freeze-substitution medium consisting of 0.01% uranyl acetate in dry acetone containing 1% H_2_O and processed at –90 °C for 9 h in a freeze-substitution unit (Leica EM AFS2, Leica Microsystems). Subsequent processing was carried out using a freeze-substitution processor (Leica EM FSP, Leica Microsystems) as follows: incubation in substitution medium at –45 °C for 14 h; washes in dry acetone at –45 °C; infiltration with HM20 methacrylate resin in dry acetone at 10%, 25%, and 50% (each for 2 h at –45 °C), 75% HM20 at –35 °C for 2 h, and 100% HM20 at –25 °C for 30 h. Resin polymerization was performed by UV illumination at –25 °C for 48 h, followed by continuous polymerization during gradual warming to 20 °C over 9 h, and an additional 24 h UV polymerization at 20 °C.

For postembedding immunogold labeling, HM20-embedded tissue was sectioned using an ultramicrotome (Leica EM ARTOS 3D, Leica Microsystems) equipped with a diamond knife. Serial ultra-thin sections (80-nm thickness, 700 µm^2^) were collected onto bare Ni400 grids. Sections were quenched with 80 mM glycine in 0.1x PBS for 15 min to neutralize residual aldehydes, followed by blocking in 5% BSA and 2% cold-water fish skin gelatin in 0.1x PBS for 30 min.

Immunolabeling was performed using mouse anti-vGlut1 IgG (1:10 dilution) and rabbit anti-LRRC57 (1A1) IgG (1:2.92 dilution) as primary antibodies, followed by 6-nm colloidal gold–conjugated donkey anti-mouse IgG (1:5 dilution) and 18-nm colloidal gold–conjugated donkey anti-rabbit IgG (1:5 dilution). All labeling steps were performed at room temperature, except for primary antibody incubation, which was done overnight at 4 °C.

Following immunolabeling, sections were coated with 7-nm carbon using a high-vacuum carbon coater (Leica EM ACE600) and imaged using a Tecnai F20 transmission electron microscope (Thermo Fisher Scientific) equipped with a 15-megapixel digital camera (NanoSprint15, AMT Imaging).

#### Electron microscopy of sections for large fields of view

Following transcardial perfusion with aldehyde fixatives (2.5% glutaraldehyde, 2% PFA, 0.1 M sucrose in 0.1 M cacodylate buffer, pH 7.4) and mannitol, hippocampal CA1 and CA3 tissue blocks (∼0.8 mm × ∼0.8 mm × ∼0.15 mm) were obtained by vibratome sectioning and manual dissection in 0.1 M cacodylate buffer.

Tissue blocks were post-fixed in 2% OsO_4_ and 1.5% K_4_Fe(CN)_6_ in cold 0.1 M cacodylate buffer for 1 h, followed by treatment with 1% thiocarbohydrazide for 20 min. A second post-fixation with 2% OsO_4_ in cold cacodylate buffer was performed to further enhance lipid membrane contrast. For *en bloc* heavy metal staining, samples were incubated in 2% uranyl acetate in 0.1 M acetate buffer overnight at 4 °C, followed by incubation in Walton’s lead aspartate solution (0.03 M L-aspartic acid, 0.66% lead nitrate, pH 5.5) for 30 min at 60 °C. Tissue blocks were dehydrated through a graded ethanol series (30%, 50%, 70%, 80%, 90%, 95%,100% (I), and 100% (II); each step was 10 min) and infiltrated with Durcupan epoxy resin for 46 h on a rotating platform. Tissue blocks were embedded in flat molds and polymerized at 70 °C for 4 days.

Resin-embedded samples were sectioned using an ultramicrotome equipped with a diamond knife. Consecutive ultra-thin (100-nm) sections spanning areas approximately 700 µm^2^ were collected onto silicon chips (5 mm × 7 mm). Sections were dried on a heat block at 45 °C for 1 h.

Electron microscopy imaging was performed using a Zeiss Gemini460 field-emission scanning electron microscope equipped with a Sense back-scattered electron (BSE) detector and Atlas 5 array tomography software. Images were acquired at 2 kV accelerating voltage, 80 pA probe current, and a 4.3-mm working distance. Regions of interest were imaged with a 45 µm^2^ field of view at 2-nm pixel resolution.

### Proteomics

Immunoprecipitated proteins were processed using SP3 bead–based digestion. Briefly, proteins were resuspended in a buffer containing 2% SDS and 50 mM HEPES (pH 8.5), reduced with 5 mM DTT at room temperature for 30 min, and alkylated with 10 mM iodoacetamide. Excess iodoacetamide was quenched by the addition of 10 mM DTT. Proteins were then bound to a 1:1 mixture of hydrophilic and hydrophobic SP3 beads by adjusting the solution to 50% ethanol and incubating with shaking at 1000 rpm for 10 min. Beads were washed 4 times with 80% ethanol and briefly air dried. Proteins were digested on-bead with trypsin at a 1:25 (enzyme:protein, w/w) ratio in 10 mM ammonium bicarbonate buffer (pH 8.5) at 37 °C overnight. Peptides were collected, acidified with formic acid, and dried prior to LC–MS/MS analysis.

For mass spectrometry analysis, dried peptides were reconstituted in 5% formic acid and separated on a reversed-phase column (75 µm × 20 cm, 1.7 µm C18 resin; CoAnn Technologies) coupled to an Orbitrap Exploris 480 mass spectrometer (Thermo Fisher Scientific) via a Dionex UltiMate 3000 nanoLC system. Peptides were eluted at 65 °C by using a 12%-36% buffer B gradient over 30 min (buffer A: 0.1% formic acid in water with 3% DMSO; buffer B: 0.1% formic acid in 67% acetonitrile with 3% DMSO) at a flow rate of 0.25 µL/min. The mass spectrometer was operated in positive-ion mode by using data-independent acquisition, consisting of one full MS scan followed by 32 MS/MS scans. MS1 spectra were acquired at a resolution of 60,000 with an AGC target of 3 × 10, a scan range of m/z 450–1100, and a maximum injection time of 25 ms. MS2 spectra were acquired at a resolution of 30,000, with a fixed first mass of m/z 120, an AGC target of 1 × 10, a maximum injection time of 22 ms, and a 20 m/z isolation window.

Raw data were analyzed using Spectronaut (version 19) against a mouse protein database containing 55,260 entries. Carbamidomethylation of cysteine was set as a fixed modification, and methionine oxidation and protein N-terminal acetylation were set as variable modifications. Peptide- and protein-level false discovery rates were controlled at 1%. Differentially expressed proteins and peptides were identified using log₂ fold-change and statistical testing implemented in the limma R package. Statistical significance was determined based on adjusted P-values and log₂ fold-change thresholds (>2 standard deviations). Protein-level standard deviations were estimated by fitting to a Gaussian distribution to assess the magnitude of experimental variation.

### LRRC57/RABIN Histology

Mice were perfused with 10% neutral buffered formalin. Heads were post-fixed for 72 h before brains were removed from the cranium and paraffin-embedded in a coronal orientation. Slides were sectioned at 4 μm for either routine H&E staining (HistoCore SPECTRA, Leica) immunohistochemistry (IHC), or immunofluorescence (IF) labeling for LRRC57. All assay steps for IHC were performed on the Bond Max with Bond wash buffer (Leica, AR9590) rinses between steps. Slides were incubated with the primary antibody 1A1 for LRRC57/RABIN at 1:50; antibody binding was detected using the anti-rabbit Bond Polymer Refine Detection kit (Leica, DS9800). IF labeling was performed on a Ventana Discovery Ultra autostainer (Roche Diagnostics). First, slides underwent heat-induced epitope retrieval (HIER) using cell-conditioning media 1 (Roche, #950-500), were incubated with antibody 1A1 for LRRC57/RABIN at 1:100, and primary antibody binding was detected with Cy5 (DISCOVERY Cy5 detection kit, Roche #760-238). HIER was performed a second time, then slides were incubated with either VGLUT1 (Synaptic Systems #135302) at 1:2000, GAD2 (Abcam #ab239372) at 1:16,000 and PCSK1 (Abcam # ab220363) at 1:1000 and detected with FAM (DISCOVERY FAM detection kit, Roche # 760-243). Slides were coverslipped using Prolong Gold antifade with DAPI. Slides were imaged on a Zeiss Axioscan slide scanner, and labeling markups were created using the Area Quantification FL v2.3.4 algorithm. Figure images were generated using HALO v3.6.4134.137.

### Inflammation and Necrosis Histology

#### Kainic acid treatment and experimental groups

Kainic acid (KA; Hello Bio, HB0355) was diluted to a concentration of 2 mg/mL in sterile injection saline and stored at –20 °C. Aliquots were thawed once and discarded after use. Mice (12–18 weeks old) were given 15-25 mg/kg KA via intraperitoneal (i.p.) injection and were monitored for 2 h postinjection. Seizure severity was scored using a modified Racine scale: Stage 1, immobility/absence-like behavior; Stage 2, hunched posture or wide stance with facial movements; Stage 3, partial rearing with forelimb clonus or tonic extension; Stage 4, full rearing with clonic, tonic, or clonic-tonic forelimb movements without loss of balance; Stage 5, similar movements with loss of balance or wild jumping; Stage 6, death^61^. At the end of the observation period or upon reaching a humane endpoint, diazepam (10 mg/kg, Patterson Veterinary, 78940890) was administered intramuscularly. Mice were singly housed for 2 days postinjection, after which females were rehoused together.

#### Perfusion and immunohistochemical staining

Mice were deeply anesthetized with tribromoethanol (6.875 mg, i.p.) and transcardially perfused with artificial cerebrospinal fluid containing (in mM) 125 NaCl, 2.5 KCl, 2 CaCl_2_, 1 MgCl_2_, 1.25 NaH_2_PO_4_, 26 NaHCO_3_, and 20 glucose, equilibrated with 95% O_2_/5% CO_2_ for 3 min at 3.5 mL/min, followed by 4% PFA in 0.1 M PB for 5 min at the same flow rate. Brains were post-fixed overnight at 4 °C in 4% PFA, washed twice in PBS with gentle rocking (25 min each), embedded in 4% low–melting point agarose (Thermo Fisher; 16520-100) in PBS, and sectioned coronally at 50-µm using a vibratome. Sections were stored in PBS containing 0.02% sodium azide.

For immunostaining, sections underwent heat-mediated antigen retrieval in 10 mM sodium citrate buffer (pH 6.0) at 80 °C for 20 min, cooled to room temperature, washed in distilled water (10 min) and PBS (20 min), and blocked for 1 h at room temperature in blocking buffer (0.02% sodium azide, 3% BSA, 5% goat serum, 0.2% Triton X-100 in PBS). Sections were incubated for 48 h at 4 °C in blocking buffer with primary antibodies against IBA1 (mouse; Synaptic Systems, 234011; 1:500– 1:1,000), GFAP (chicken; Synaptic Systems, 173006; 1:500–1:1,000), and cleaved caspase-3 (rabbit; Cell Signaling Technology; 9661S at 1:500). After four washes in PBS containing 0.1% Tween-20 (PBS-T; 20 min each), sections were incubated with species-specific fluorophore-conjugated secondary antibodies (1:1000; Invitrogen A32931, Invitrogen A11031, Biotium custom CF dye, lot 23C0331) in blocking buffer for 48 h at 4 °C. Sections were washed twice in PBS-T (20 min each), counterstained with DAPI (1:1000 in PBS; Invitrogen) for 30 min at room temperature, washed twice in PBS (20 min each), mounted with ProLong Gold antifade mounting medium (Invitrogen, P36935), and imaged by confocal microscopy (20×, 0.8 NA air objective for GFAP and Iba1; 10×, 0.45 NA air objective for cleaved caspase-3) using a Nikon AX microscope.

#### Quantification (general)

For EEG-recorded animals, only the hemisphere contralateral to the electrode implantation was analyzed; for all other animals, both hemispheres were analyzed. Values were averaged across hemispheres and across three sections per mouse to generate a single value for normalization and statistical analysis. All quantification was performed blinded to genotype.

#### Quantification (hippocampus)

Z-stack images of the CA1 region were acquired at 20× magnification (z-step, 0.63 µm) by using a 3i Marianas W1 microscope. Maximum-intensity projections of 10 consecutive z-planes were generated in Fiji (ImageJ). Regions of interest (ROIs) encompassing the pyramidal cell layer were manually delineated; astrocytes and microglia were identified by morphology and were manually counted using the Cell Counter plugin in Fiji. Cell density was calculated as cells per ROI.

#### Quantification (cortex)

Two adjacent cortical fields (immediately dorsal to the hippocampus and directly dorsal to the first field) were imaged at 20× magnification by using identical settings and stitched using Pairwise Stitching in Fiji (Preibisch et al., 2009). A standardized 200-µm-wide ROI spanning layers I–VI of the cortex was applied. IBA1- and GFAP-positive cells were manually counted using the Cell Counter plugin, and cell density was calculated per ROI.

#### Quantification (cleaved caspase-3)

Cleaved caspase-3^+^ cells in the hippocampus and cortex were quantified from images acquired using a Nikon AX microscope at 10× magnification. Z-stack images were collected from 10-µm-thick tissue sections with five optical planes spaced 3-µm apart. Three serial sections per mouse were analyzed. Positive cells were identified based on morphology and fluorescence-intensity threshold and were manually counted using the Cell Counter plugin in Fiji. Counts were averaged across sections for each mouse and were used for statistical analysis.

### Single-Cell Electrophysiology

#### Brain slice preparation

Mouse brains were removed and placed in cold (4 °C) dissecting media containing (in mM) 125 choline-Cl, 2.5 KCl, 0.4 CaCl_2_, 6 MgCl_2_, 1.25 NaH_2_PO_4_, 26 NaHCO_3_, and 20 glucose (300–310 mOsm), equilibrated with 95% O_2_/5% CO_2_. Coronal slices (350 μm) containing dorsal hippocampus and somatosensory cortex were made using a vibrating microtome (Leica VT1200). Slices were transferred to artificial cerebrospinal fluid (ACSF) containing (in mM) 125 NaCl, 2.5 KCl, 2 CaCl_2_, 2 MgCl_2_, 1.25 NaH_2_PO_4_, 26 NaHCO_3_, 20 glucose (300–310 mOsm), equilibrated with 95% O_2_/5% CO_2_ at 34 °C for 30 min, followed by 1 h at room temperature prior to use. Slices were transferred to a recording chamber mounted on an upright microscope (Olympus BX51WI) and superfused (1-2 mL/min) with warm (30-32 °C) ACSF. Slices were viewed with a CCD camera (Rolera-XR, QImaging) using IR-DIC optics. CA1 pyramidal neurons were identified by soma shape, size, and location within the pyramidal cell layer. Cortical layer 2/3 cortical pyramidal neurons were identified by soma size and shape.

#### Whole-cell recording

Whole-cell recordings were made with patch pipettes (3-6 MOhm) using a Multiclamp 700B amplifier, digitized (10 kHz) with a Digidata 1440, and recorded using pCLAMP 10 software (all Molecular Devices). In all experiments, membrane potentials were corrected for a liquid junction potential of –10 mV. In voltage-clamp recordings, series resistance, input resistance, and holding current were monitored for stability. Cells with series resistance more than 30 MOhms or cells that changed resistance values more than 20% over the duration of recordings were rejected. Drugs were added to ACSF.

For standard voltage-clamp recordings, patch pipettes were filled with an internal solution containing (in mM) 125 CsMeSO_3_, 2 CsCl, 10 HEPES, 0.1 EGTA, 4 ATP-Mg_2_, 0.3 GTP-Na, 10 creatine phosphate-Na_2_, 5 QX-314, and 5 TEA-Cl (pH 7.4, 290-295 mOsm).

Miniature spontaneous synaptic inputs were recorded in the presence of 0.5 µM tetrodotoxin (TTX). For excitatory postsynaptic currents neurons were held at –70 mV with inhibitory inputs blocked by 100 µM picrotoxin. For inhibitory synaptic currents neurons were held at 0 mV with excitatory inputs blocked with 3 mM KA. Spontaneous activity was recorded for 5–10 min, beginning at least 2 min after whole-cell break-in. Spontaneous postsynaptic currents were detected using miniAnalysis (Synaptosoft), as deviations of more than five times the baseline root-mean-squared noise level.

Miniature excitatory postsynaptic currents were recorded from cultured hippocampal neurons (preparation described below) as in slices. Likely pyramidal neurons were targeted based on large, pyramidal cell bodies with thick primary dendrites.

### Primary Culture of Mouse Hippocampal Neurons

Primary hippocampal neuron cultures were prepared from E18–E20 C57Bl/6J mouse embryos. Pregnant dams were euthanized by rapid decapitation, and embryos were removed from the uterine horns and immediately decapitated into ice-cold dissection solution (DS) consisting of 1× HBSS (Thermo Fisher; 14185052) supplemented with 1 mM sodium pyruvate (11360070), 10 mM HEPES (15630080), penicillin-streptomycin (15140122), and 30 mM glucose (Sigma; G7528).

After removal of overlying skin and skull with fine forceps, brains were extracted and hemisected to separate the cerebral cortex from the hindbrain. The hippocampi were carefully dissected from the neocortex and transferred to fresh ice-cold DS. Once both hippocampi were collected, the DS was replaced with room-temperature papain digestion solution, consisting of papain (Worthington; LK0031761) and DNase I (Worthington; LK003170) dissolved in DS. Tissue was incubated for 30 min at room temperature with gentle rocking. Then it was washed in pre-warmed plating medium, which consisted of Neurobasal Plus, (Thermo Fisher; A3582901) supplemented with 2% B27 Plus (Thermo Fisher; A3582801), 2% GlutaMAX (Thermo Fisher; 35050061), 1% fetal bovine serum (Hyclone; SH30071.03), and 5 µg/mL Primocin (InvivoGen; ant-pm-1). Tissue was gently dissociated by trituration using a 5-mL plastic serological pipette, and cells were resuspended in platin medium. Cell density was determined by trypan blue (Thermo Fisher; 15250061) exclusion using a hemocytometer. Cells were plated into tissue culture plates treated with either 1.25 × 10^5^ (12-well, Corning; 3513) or 5 × 10^5^ cells (6-well, Corning; 3516). For fluorescence microscopy, neurons were grown on 18-mm #1.5 German glass coverslips in 12-well plates (Warner Instruments; CS-18R15). UV-sterilized coverslips were incubated overnight in 0.1 mg/mL poly-D-lysine (Sigma, P7280) dissolved in 22-μm filter-sterilized borate buffer (50 mM boric acid, 12.5 mM sodium borate, pH 8.5). The following day, coverslips were washed using sterile water and coated with 0.01 mg/mL mouse laminin (ThermoFisher, 23017015) in PBS for more than 2 h. After seeding, plating medium was replaced with Neurobasal Plus Medium (Invitrogen) supplemented with 2% B27 Plus and 1% Glutamax. Neurons were fed 2-3 times per week thereafter.

### Fluorescence Imaging and Field Stimulation of Hippocampal Neurons

Fluorescence imaging was performed on an Olympus IX83 inverted microscope equipped with a 100×/1.5 NA oil-immersion objective (UPLAP0100XOHR), an X-Cite XYLIS LED light source, and an ORCA-Fusion BT sCMOS camera (Hammamatsu Photonics). Primary hippocampal neurons were transduced with lentivirus between days in vitro (DIV)2 and DIV4, and imaging experiments were conducted between DIV15 and DIV24.

For imaging, neurons were transferred to a field-stimulation chamber (Warner Instruments; RC-49MFSH), and culture medium was exchanged with the following extracellular solution (in mM unless otherwise stated): 135 NaCl, 5 KCl, 5.5 glucose, 25 HEPES, 2 CaCl_2_, 1 MgCl_2_, supplemented with 2% B27 plus, 1% GlutaMAX (pH 7.3). Temperature (33-35 °C) were maintained with an Okolab environmental controller and with a stage-top incubation chamber.

Fluorescence excitation was delivered using a 490/20-nm bandpass filter, and emission was collected through a 525/36-nm bandpass filter, thus capturing single focal planes. Field stimulation was delivered using a Model 4100 Isolated High-Power Stimulator (A-M Systems; 930000) and synchronized with image acquisition via an Axon Digidata 1550B (Molecular Devices). Field-stimulus voltage was set to the minimum level that reliably evoked presynaptic calcium transients in >95% of boutons, which was determined using neurons transduced with synaptophysin-GCaMP6f under the control of the hSynapsin promoter (pFUG-hSyn-vGlut1-pHluorin).

### iGluSnFr Imaging

Primary neurons were transduced with a lentiviral vector encoding iGluSnFr3.v857 under control of the mouse CaMKIIα promoter (pFUG-mCaMKIIα-iGluSnFR3.v857). For spontaneous glutamate release imaging, action potentials were blocked by adding 1 µM tetrodotoxin citrate (Tocris; 1069) to the extracellular solution. Time series were acquired at 20 Hz for 60 s (1200 frames) at 65 nm per pixel resolution. All synapses exhibiting activity during the acquisition period were manually segmented in ImageJ, and the fluorescence-intensity traces were exported for analysis of event frequency and amplitude using a custom Microsoft Excel–based algorithm.

For field stimulation-evoked iGluSnfr experiments, postsynaptic activity was blocked by supplementing the extracellular solution with (in µM) 50 D-AP5 (HelloBio; HB0025), 20 CNQX (HelloBio; HB0205), and 100 picrotoxin (Tocris; 1128). Time sequences were acquired at 100 Hz for 2.0 s (200 frames). After 90 baseline frames, a field stimulus was delivered, followed by a second stimulus after 200, 500, or 1000 ms. The paired-pulse ratio was calculated as the peak amplitude of the second response divided by that of the first.

For time-to-peak analysis, active synapses were segmented using a custom semiautomated ImageJ macro, and fluorescence traces were exported to IgorPro (Wavemetrics) for kinetic analysis.

### vGlut1-pHluorin Imaging

Primary neurons were transduced with a lentiviral vector encoding vGlut1-pHluorin under control of the human Synapsin (hSyn) promoter (pFUG-hSyn-vGlut1-pHluorin). During imaging, cultures were perfused with extracellular solution supplemented with (in µM) 50 D-AP5 (HelloBio; HB0025), 20 CNQX (HelloBio; HB0205), 100 picrotoxin (Tocris; 1128) to block postsynaptic activity.

Synaptic vesicle pool sizes were quantified at individual synapses based on stimulus-evoked changes in vGlut1-pHluorin fluorescence as previously described^62^. Two identical 100 Hz, 50-spike field stimuli designed to evoke exocytosis of the readily releasable pool of synaptic vesicles. The first stimulus allowed for measurements of vesicle recycling and endocytosis rate. The second stimulus was carried out after wash-in of 80 nM folimycin (Tocris; 2656), a specific inhibitor of the vacuolar-type ATPase (V-ATPase) to prevent vesicular re-acidification. Previously exocytic synaptic vesicles remain stably fluorescent after endocytosis to estimate the size of the exocytic vesicle pool. Next, a sustained 40 Hz, 900-spike field stimulation depletes the reserve vesicle pool. Finally, the alkylation of total vGlut1-pHluorin by superfusion of imaging solution adjusted for osmolarity with the NH_4_Cl (in mM: 85 NaCl, 50 NH_4_Cl, 5 KCl, 5.5 glucose, 25 HEPES, pH 7.3) allowed for estimating the nonexocytic resting pool.

### Mouse Behavioral Assays

#### Open Field

Mice were placed in a 41 cm square opaque plastic arena and allowed to explore for 15 min. Locomotion (distance traveled) was recorded using AnyMaze (Stoelting, Illinois) with center-of-mass tracking. *Elevated Plus Maze.* Mice were placed on a plus-shaped arena (30 x 6.4 cm arms, 61 cm elevated). Behavior was recorded and analyzed using TopScan (CleverSys, Reston, Virgina) with center-of-mass tracking. Open-arm exploration was defined by tail-base entry and expressed as % distance in the open arm = (open-arm distance / total distance) x 100. *Social Preference.* Mice were placed in a 41 cm square opaque arena and allowed to explore for 5 min. Tracking and scoring were performed using AnyMaze (Stoelting, Illinois). Exploration was defined as head position within 2.5 cm of either stimulus. Social preference was calculated as (time exploring mouse)/(time exploring mouse + object) x 100.

### Electroencephalography and Electromyography

#### Telemetry unit implantation and data collection

Electroencephalograms (EEG) and electromyograms (EMG) in mice were recorded using a wireless telemetry system (Ponemah v6.51, Data Systems International). Mice (12–16-weeks old), weighing at least 21 g, were implanted with a telemetry unit (HD-X02), according to the manufacturer’s instructions. Briefly, mice were injected with extended-release meloxicam preoperatively, then anesthetized using isoflourane vapors (5% induction, 2% maintenance). After the loss of the pedal reflex, fur was removed from the head and the left flank, and the surgical area was prepped. The animal was then placed into the stereotaxic device and ophthalmic ointment was applied. A midline incision was made at the posterior margin of the eyes extending 2-3 cm. A subcutaneous pocket was made along the dorsal flank, into which the unit was placed with the leads projecting rostrally. Two leads were then secured to the cervical trapezius muscle. Two fenestrae were made in the skull, +2 mm and –2 mm from bregma, –1.5 mm and +2 mm from midline, respectively. A screw was placed into each hole, an EEG lead was wrapped around each screw, and then the screws were tightened, such that the screw tip touched dura. Dental acrylic was used to secure the leads in position and electrically isolate the leads from surrounding tissue. The incision was then sutured closed (Vicryl) and wound clips were applied. Mice were then placed in a clean cage on a heating pad to recover until the onset of data collection. All procedures were conducted using aseptic techniques with sterile instruments. Mice were individually housed postoperatively and for the EEG/EMG recording.

Continuous recording started 7–10 days postoperatively, and sampling frequency was at 500 Hz. Baseline data were collected for 2–4 days before i.p. KA injection, subthreshold (10 mg/kg) or threshold dose of 20 mg/kg (females) and 25 mg/kg (males), both doses based on in-lab pilot studies. KA was delivered 2–3 h after light onset. Data were collected for as many as 4 days post-treatment. Video was recorded for each animal for the entirety of the EEG/EMG recording.

#### EEG and EMG Data Analysis

Data were analyzed offline by using a software-based, semiautomated method (Neuroscore v3.6.0, Data Science International). The signals were processed using the Hamming window function in 10-s epochs. Seizure-like activity was defined by high amplitude activity (>2× the background), with a frequency >5 Hz and a minimum duration of 10 s^63^. The spike trains identified were then visually verified by a trained observer using the waveform and video to eliminate artifacts and identify and score spontaneous and KA-induced seizures and seizure-like activity by using a modified Racine scale^61^. To quantify spiking events/spike wave discharges, the same parameters (as mentioned above) were used to identify spikes, apart from raising the threshold ratio to 1:7. All flagged individual spikes were then visually verified and artifacts were excluded.

### Statistics and Reproducibility

No statistical methods were used to predetermine sample size. Sample sizes were based on previous studies using similar methodologies. Mice of both sexes were used throughout. Animals were assigned according to breeding outcome, and littermate controls were used whenever possible. Investigators were blinded to genotypes or treatments during data analysis whenever feasible.

The exact sample size (n) and unit of measurement (e.g., mice, cells, cultures, synapses, or fields of view [FOVs]) for each experiment are provided in the figure legends. Measurements were obtained from distinct biological samples unless otherwise indicated. For imaging experiments, synaptic measurements were averaged within each FOV before statistical comparison. Data are presented as mean ± s.e.m. unless otherwise indicated.

Normality was assessed before parametric testing. Two-group comparisons were performed using two-sided Welch’s t-tests or Mann–Whitney U tests, as appropriate. Repeated-measures and multi-factor experiments were analyzed using two-way ANOVA followed by appropriate post hoc multiple-comparison testing. No covariates were included in the statistical analyses unless explicitly stated. Exact statistical tests, sample sizes, test statistics, degrees of freedom (where applicable), and exact *P* values are provided in the figure legends.

Animals exhibiting extensive hippocampal necrosis that precluded accurate quantification of inflammatory or apoptotic markers were excluded from those specific histological analyses according to predefined criteria. No other samples were excluded. Key findings were independently replicated across multiple animals, cultures, and experimental approaches, including electrophysiology, optical imaging, electron microscopy, biochemistry, proteomics, behavior, and EEG recordings. Statistical analyses were performed using GraphPad Prism and associated analysis software.

